# Cell migration sculpts evolutionary dynamics favouring therapy resistance in lung cancer

**DOI:** 10.1101/2025.09.15.675999

**Authors:** Ajay Bhargava, Xiao Fu, Sasha Bailey, Yutaka Naito, Hamid Mohammadi, Robert Hynds, Steven Hooper, Dhruva Biswas, Alix Le Marois, Sunil Kumar, Yuriy Alexandrov, Paul French, James McGinty, Paul A. Bates, Charles Swanton, Erik Sahai

## Abstract

The evolution of cancer undermines the long-term efficacy of therapy. In this study, we combine experimental, computational, and clinical data analysis to investigate factors influencing the competition of subclones in lung cancer. Lineage tracing reveals unexpected variation in the long-term fate of neutral subclones, with subclones arising near the edge of tumours being favoured. Low levels of cell migration and cell mixing lead to high cell densities in the interior of the tumour that suppress proliferation. Using agent-based modelling and *in silico* analysis we inferred the extent of cell mixing in human tumours from the TRACERx lung cancer study. This reveals correlations between Epithelial to Mesenchymal Transition (EMT), stromal fibroblasts and levels of cell mixing. Experimental analysis confirms that both TGFβ-driven EMT and stromal fibroblasts reduce the variability in subclone fate and promote subclone mixing. Moreover, mixing favours clonal sweeps by subclones resistant to therapy-induced cell killing. Together, these analyses demonstrate that EMT and stromal fibroblasts sculpt tumour evolution by promoting cell mixing and thereby favour the rapid dominance of therapy resistant subclones.

## Introduction

Cancer arises as the result of mutations within somatic cells that confer some selective advantage (Nowell, 1976). Ultimately, this enables cancer cells to break free from the regulatory mechanisms governing normal tissue homeostasis. This evolutionary process presents a challenge for both understating the basic mechanisms of tumour progression and cancer therapies. Over time, cells containing mutations or chromosomal alterations conferring advantageous phenotypes are selected for, leading to more aggressive tumour phenotypes. Understanding the long-term fate of an individual cancer cell’s progeny has the potential to reveal how the interplay between cell intrinsic factors and the extrinsic cellular context shape evolution. In experimental models, this tracking can be achieved by using irreversible Cre or Flippase recombinase-based labelling methods (Orban, Chui and Marth, 1992). Several recent studies have exploited this technology in models of skin and intestinal tumorigenesis (Lamprecht *et al*., 2017; Lenos *et al*., 2018; Reeves *et al*., 2018; Van Der Heijden *et al*., 2019).These tissues are characterised by actively proliferating stem cells in their normal state and tumour studies have revealed that this situation persists in cancer, albeit with some dysregulation. Mutations arising in stem cells have a greater propensity to persist and expand, with the population of cells containing the mutation termed a clonal expansion. In contrast, mutations arising in cells committed to terminally differentiate are unlikely to persist. Tumours arise as the result of a clonal expansion in a tissue. However, they are not genetically uniform and contain multiple subclones, some of which may have varying fitness. Extrinsic factors can also influence the likelihood of mutations persisting; in intestinal tumours stromal fibroblasts contribute to defining a stem cell niche at the tumour edge and favour the expansion of subclones arising in this location (Lenos *et al*., 2018). The situation in tumours that arise from epithelial tissues that largely quiescent, such as the lung, is much less clear.

In addition to experimental models, several computational studies investigated the fate of cells and evolutionary dynamics in diverse multi-cellular systems spanning bacterial biofilms to cancer. These studies highlighted the role of spatial context (Fusco *et al*., 2016; Chkhaidze *et al*., 2019; West *et al*., 2021; Fu *et al*., 2022; Noble *et al*., 2022), physical cell-cell interactions (Farrell *et al*., 2017; Kayser *et al*., 2018), and cell dispersal (Waclaw *et al*., 2015; Paulose and Hallatschek, 2020) in determining the fate of subclones. These studies introduced additional concepts, suggesting that spatial position or migratory dynamics might influence clonal dynamics. Nevertheless, the molecular and cellular mechanisms influencing competition of subclones within a tumour and whether such mechanisms can be observed in patients remains unclear. This is partly due to the low temporal frequency of obtaining patient tissue and challenges in determining subclonal relationships and spatial organisation with high accuracy.

Therapeutic regimens apply a strong selective pressure, with the expansion of rare pre-existing or de novo arising subclones of cells resistant to the therapy a frequent outcome (Piotrowska *et al*., 2015; Hata *et al*., 2016; Quek *et al*., 2018). This is termed ‘acquired resistance’ and often linked to increased metastatic spread with frequently fatal consequences. While predicting evolutionary trajectories is problematic, understanding of the mechanisms of competition between cells of differing fitness will lead to improved therapeutic strategies. In particular, understanding the ‘rulebook of cancer evolution’ should lead to cancer therapies that sculpt tumour evolution towards less aggressive and less life-threatening states.

By combining experimental and computational approaches, we reveal that the fate of subclones in lung cancer is highly variable – which we term a ‘place of birth’ effect. This variability arises from low levels of cell migration leading to increased compressive stress and exit from the cell cycle. Agent-based modelling of different migratory regimes and their associated subclonal dynamics is used to infer the ‘mixedness’ of human lung adenocarcinoma. This analysis confirms that promoting cell migration, either via TGFβ-driven EMT or by stromal fibroblasts, counteracts the place of birth effect. Ultimately, increased migration and mixing enhances the rate at which drug resistant subclones become dominant. Thus, we relate cellular level migration behaviours to tumour evolution.

## Results

### Lineage tracing reveals non-equivalent fates of neutral subclones

To investigate cancer cell clonal dynamics in lung cancer, we implemented a fluorescent reporter lineage tracing approach in a KRAS and TP53 mutant murine model of lung adenocarcinoma (a C57BL/6 KRasG12D; Trp53 fl/fl cell line termed KP)(Jackson *et al*., 2005). These cells constitutively express tamoxifen regulated Cre recombinase (CreERT2), and were engineered to contain a modified ‘brainbow’ cassette (Livet *et al*., 2007; Loulier *et al*., 2014) that would recombine following the administration of tamoxifen (Figure 1a & Supp. Figure 1a&b). Each ‘position’ of the cassette contained a different fluorescent protein that could be tracked, with the progeny of each labelling event termed a subclone. Analysis of growth rates confirmed that their fates were equivalent regardless of the fluorophore expressed (Supp. Figure 1c). Thus, the subclonal labelling is neutral in evolutionary terms. Figure 1b shows tumours seeded via intravenous injection and subclonally labelled by tamoxifen injection three days later. Intriguingly, the subclones had widely varying sizes despite being induced at the same time. Optical projection tomography and vibratome sectioning both suggested that larger subclones were near the tumour surface (Figure 1b&c and Movie 1). Quantitative analysis revealed an inverse relationship between subclone size and subclone frequency, which is not consistent with a simple model of all cells having equivalent fate (Figure 1d). The variation in subclone fate was also reflected in the high standard deviation of subclone sizes (Supp. Figure 1e). This metric was high when subclonal labelling was performed early (labelled in blue), but declined if labelling was performed later (labelled in green or purple) as there was less time for differences in subclone fate to be manifested (Supp. Figures 1e&f).

**Figure 1:**
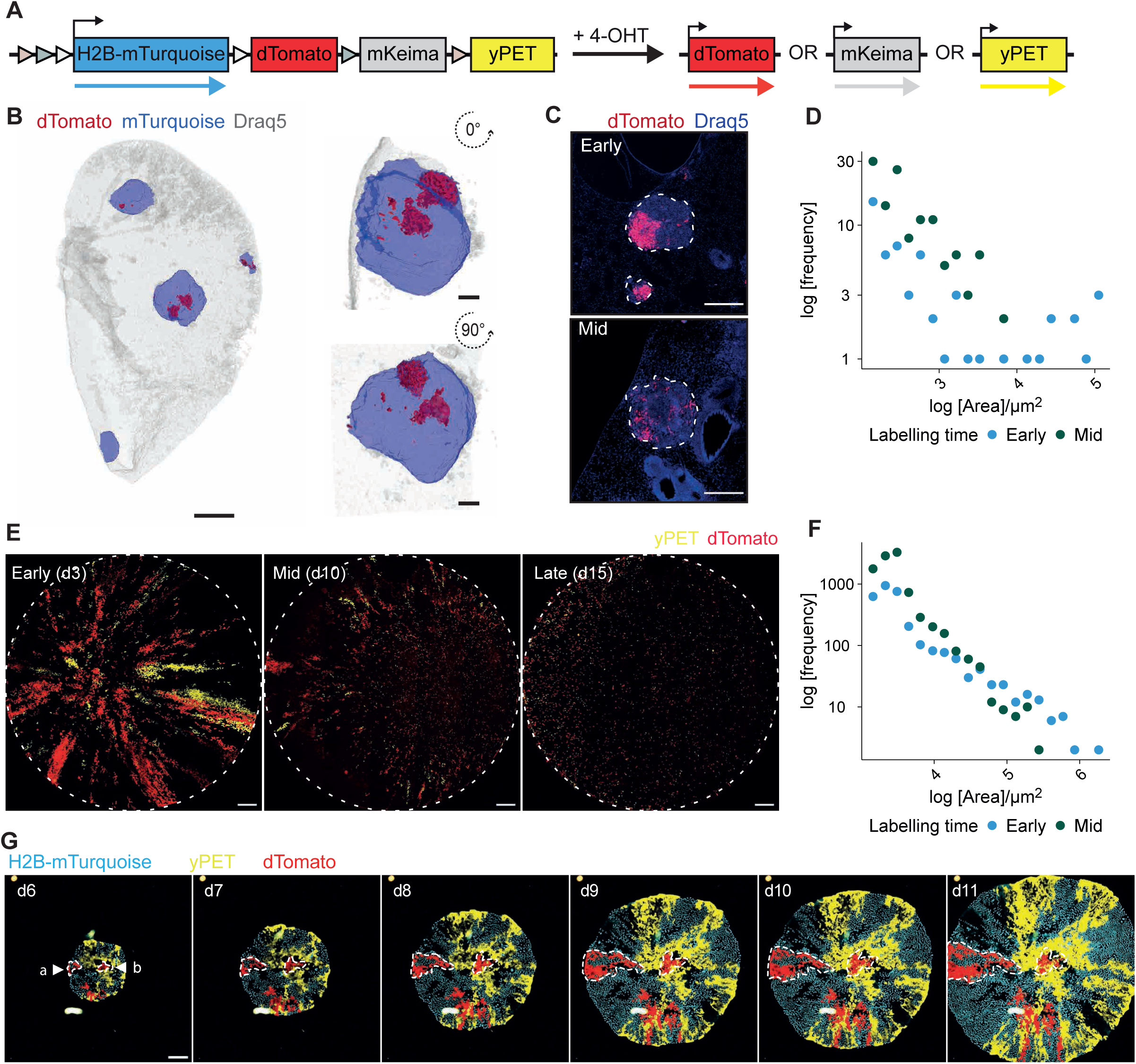
Multi-colour lineage tracing reveals unequal expansion of neutral subclones. A. Schematic of the pOncobow cassette. Unrecombined, H2B-mTurquoise is expressed, then upon 4-hydroxytamoxifen (4-OHT) treatment, dose-dependent stochastic Cre-mediated recombination induces dTomato, yPET or mKeima fluorescent protein expression for lineage tracing. B. Optical projection tomography digital reconstruction of *ex vivo* lung KP-pOncobow tumours harvested from Prkc^Skid^ mice on day 26 post-injection following 4-OHT treatment on day three of growth. mTurquoise and dTomato are shown for whole lung scale bar = 1000 µm and zoom of tumour at two rotations scale bar = 100 µm. 2x10^4^ cells were injected into the tail vein of Prkc^Skid^ mice, 20 µg g^-1^ 4-OHT was administered on day 3 (early) or day 10 (mid) post-injection, lungs were harvested on day 26 and imaged in 3D using optical projection tomography, N = 6 lungs per labelling time from two mouse cohorts. C. Confocal images of vibratome sectioned KP-pOncobow tumours from early (day 3 post-seeding) and mid (day 10 post-seeding) labelling time points. dTomato subclones and Draq5 staining are shown dotted lines denote tumour boundary, scale bars = 500 µm. D. Frequency distribution of log_10_ subclone area quantified from 2D confocal images of KP-pOncobow tumours labelled at early or mid time points. Subclone area was defined as contiguous single colour patches greater than a minimum cell area of 100 µm^2^. N = 6 lungs per labelling time from two mouse cohorts. E. Representative images of subclone patterns in single cell colonies labelled at different timepoints. Following single cell plating, cells were treated with 1 nM 4-OHT for 6 h on day 3 (early), 10 (mid) or 15 (late), then colonies were fixed on day 21 of growth. dTomato and yPET subclones are shown, scale bar = 500 µm. F. Frequency distribution of log_10_ subclone area quantified from KP-pOncobow colonies labelled at early or mid time points. Subclone area was defined as contiguous single colour patches greater than a minimum cell area of 100 µm^2^. Early n = 14, mid n = 13 colonies per group for 2 two independent experiments. G. Representative time lapse images of KP-pOncobow colony expansion every 24 h starting on day 6 of colony growth until day 11. Colours represent subclones, H2B-mTurquoise (cyan), yPET (yellow), dTomato (red). Scale bar = 500 μm, arrows denote a: edge subclone, b: centre subclone.

To narrow down the cause of the unequal subclonal fates, we sought to simplify the system *in vitro.* KP lung cancer colonies grown either as spheroids in soft agar or simply from single cells in cell cultures dishes also exhibited subclones of widely differing sizes (Figure 1e and Supp. Figure 1d), with an inverse relationship between subclone size and frequency observed once again. Of note, large wedge-shaped subclones and small round subclones were observed. Similar to the *in vivo* observations, large subclones were typically near to the tumour colony boundary, with the standard deviation of subclone size reducing the later that subclonal labelling was triggered (Figure 1e-g). Analysis of subclone size related to subclone position confirmed that subclones at or near the colony boundary exhibited the largest sizes and greatest variability in size (Supp. Figure 1g). The size of subclones in the interior was similar regardless of the timing of subclonal labelling (Supp. Figure 1g). Sequential daily imaging of colonies revealed that subclones at the colony edge expanded as streaks or wedges, whereas those in the interior showed minimal expansion (compare ‘a’ and ‘b’ in Figure 1g). Thus, we observe variability in subclone fate depending on its ‘place of birth’, with subclones initiating near the colony boundary being favoured.

The expansion of subclones at the colony edge could potentially be explained if cells nearer the edge were more proliferative, to explore this idea we engineered KP lung cancer cells to express the FUCCI cell cycle reporter (Koh *et al*., 2017). This revealed that while colonies were uniformly proliferative when small, there was a transition to a non-proliferative state in the colony interior as they became larger (Figure 2a). A similar pattern of proliferation was observed in KP tumours growing in the lung (Figure 2b & Supplementary Figure 2e). Of note, the thickness of the proliferative zone is considerably larger than that of a cell, indicating that the suppression of cell proliferation is not simply linked to the formation of cell-cell contacts on all sides (Eagle and Levine, 1967; Martz and Steinberg, 1972)(Supplementary Figure 2a). Two lines of evidence indicated that high cell density may be linked to reduced proliferation in the tumour colony interior (Figure 2c). First, we noted a greatly reduced nuclear area of cells in G0/G1 in the colony interior. This could not be explained by the reduced DNA content of G0/G1 cells as only a small difference in size was observed in small colonies that had uniform proliferation or in G0/G1 cells near the colony boundary (Figure 2c-e). Second, if tumour colonies grew into the rigid sides of the cell culture well, then both high cell density and proliferative arrest were observed (Supplementary Figure 2b&c). To determine if high cell density was a causal factor in the proliferative arrest, we removed a patch of cells in the interior of colony that had undergone proliferative arrest, thereby generating space and relieving the cell crowding. Figure 2f shows that previously arrested subclones now expanded into the space provided (see also Supplementary Figure 2d). We also tested whether seeding cells at different densities would influence their ability to proliferate. Figure 2g shows that high cell density did indeed reduce the proportion of cells in S-phase (see also Supp. Figure 2f). Thus, the high cell density observed in the centre of tumour colonies is sufficient to induce proliferative arrest. We next tested if human lung cancer cell lines would generate colonies with high densities of non-proliferative cells in their centre. We observed a range of colony phenotypes. HCC827 cells behaved in a similar manner to KP lung cells (Figure 2h). A549 and PC9 both exhibited increased cell density in the colony interior, but the reduction in cell proliferation was less apparent. Nonetheless, when normalised to DAPI intensity, the level of EdU staining in the colony centre was lower in all cell lines (Figure 2i).

**Figure 2:**
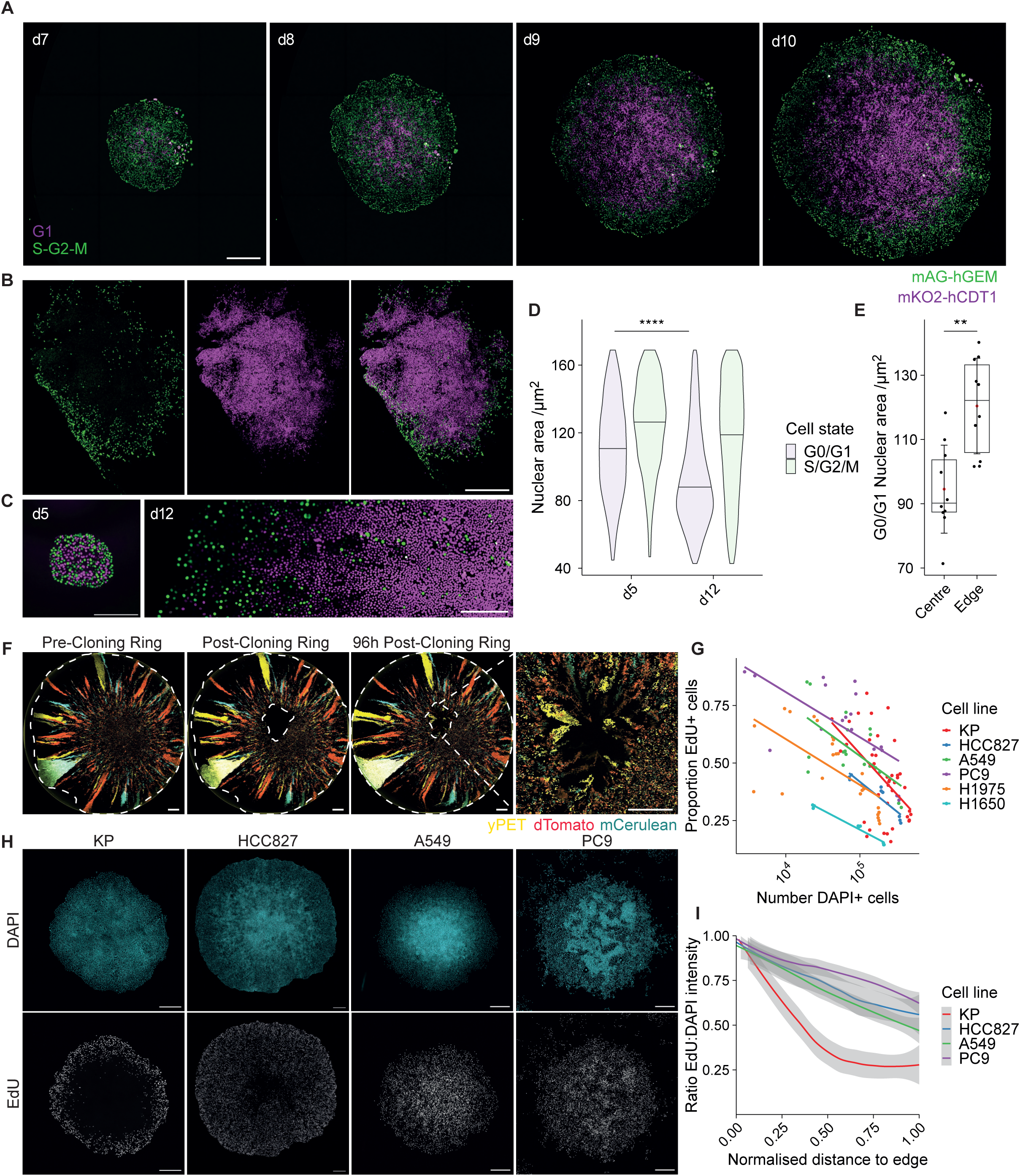
Analysis of spatial growth patterns. A. Representative time lapse images of KP-FUCCI single cell colony expansion every 24h starting on day 7 of colony growth until day 10. Geminin-Azami Green (mAG-hGEM) is shown in green and Cdt1-Kuabira-Orange (mKO-CDT1) is shown in magenta, N = 2 independent experiments, n = 4 colonies, scale bar = 500 μm. B. Representative image of KP-FUCCI tumour in an *ex vivo* lung slice from a NSG mouse. Geminin-Azami Green (mAG-hGEM) is shown in green and Cdt1-Kuabira-Orange (mKO-CDT1) is shown in magenta. N = 2 independent experiments, n = 8 mice, scale bar = 500 μm. C. Representative images of KP-FUCCI colonies at day 5 and 12 of growth. Scale bar = 500 μm D. Quantification of nuclear area following StarDist segmentation grouped by cell state, N = 3 independent experiments, day 5 n = 874 nuclei from 5 colonies, day 12 n =58,440 nuclei from 10 colonies. Median is shown as horizontal line, pairwise comparisons by Wilcoxon signed rank test with Benjamin-Hochberg p-value correction, **** p < 0.0001. E. Quantification of Cdt1-KO positive G0/G1 nuclear area for day 12 colonies following StarDist segmentation grouped by nuclear position relative to colony periphery. Regions were defined by subclone normalised minimum distance to colony boundary: edge < 0.15% and centre > 0.85% along colony radius from periphery. Error bars are mean ± sd, pairwise comparisons by Wilcoxon signed rank test with Benjamin-Hochberg p-value correction, ** p < 0.01, N = 3 independent experiments, n = 10 colonies. F. Representative images of KP-pMAGIC cytobow single cell colony grown to the area of the well (pre-cloning ring) then immediately after a cloning ring was used to remove of a patch of cells at the colony centre (post-cloning ring) and after 4 days of growth (4 days post-cloning ring). Inset is zoom of central region 4 days post cloning ring removal. Subclonal labelling was induced with 4-OHT on day 3 of growth mCerulean, dTomato and yPET subclones shown, white dashed lines denote colony periphery and region removed with cloning ring. Scale bar = 1000 µm, N = 3 replicates. G. Quantification of proportion of EdU positive cells for each cell line against total cell number at end point. KP N = 6, HCC827, A549, PC9, H1975, H1650 N = 3. H. Representative images of KP, HCC827, A549 and PC9 single cell colonies endpoint. A 1 h 10 μM EdU pulse was performed prior to fixation and immunostaining for DAPI and EdU. Scale bar = 500 μm I. Quantification of the ratio of EdU:DAPI signal intensity for 500 randomly sampled 76 x 76 μm square regions scaled to maximum colony intensity signal and plotted against region position relative to normalised distance to the colony boundary. KP n = 12, HCC827 n = 9, A549 n = 24, PC9 n = 12 colonies per condition, N = 3 independent experiments.

We speculated that the mechanical context might differ between the colony interior, colony edge, and also colony edges that ‘collided’ with a barrier. Building upon previous insights, we hypothesized that the colony interior might be under compressive stress as a result of forces generated as the colony expanded (Delarue *et al*., 2014; Streichan *et al*., 2014; Irvine and Shraiman, 2017; Di Meglio *et al*., 2022). To test if forces generated by an expanding colony would push their surroundings or on cells in the colony interior, we established an assay in which an unbounded KP cells met a deformable boundary. These analyses revealed that KP cells compress the boundary (Supp. Figure 3a), confirming their ability to generate pushing forces. Together with the increased cell density and smaller nuclear size (Figure 2a-e), these data support the hypothesis that the centre of colonies are under compressive stress.

### Migratory dynamics are linked to proliferative patterns

To understand the causes of non-equivalent sizes of subclones and the varying behaviour of different cell lines we performed time-lapse imaging. This confirmed the radial expansion of subclones located near the colony edge that was inferred from longitudinal imaging (Figure 3a-c Movie 2&3 also c.f. Figure 1h). Cells near the colony edge moved outwards with high migration persistence values. In contrast, cells in the interior showed very low levels of cell migration and low persistence (Figure 3b&c). Analysis of lung slice cultures of KP tumours grown *in vivo* confirmed the increased persistence of cancer cells at the tumour edge (Supp. Figure 3a&b). Together, these results suggest a linkage between low cell migration, high cell density, and exit from the cell cycle in the KP lung model. As detailed in Figure 2, A549, HCC827, and PC9 cells generate colonies with different patterns of cell density and proliferation. Intriguingly, cell migration analysis of these models revealed that the emergence of a non-proliferative interior zone was inversely correlated with the migratory capability of the different human cancer cells, with the most migratory PC9 cells yielding the smallest decrease in proliferation in the colony interior (Figure 3d, see also Figure 2h&i). These data suggest that high cell migration precludes the build-up of a dense non-proliferative zone and, therefore, will reduce the variability of subclone fate and modify evolutionary dynamics.

**Figure 3:**
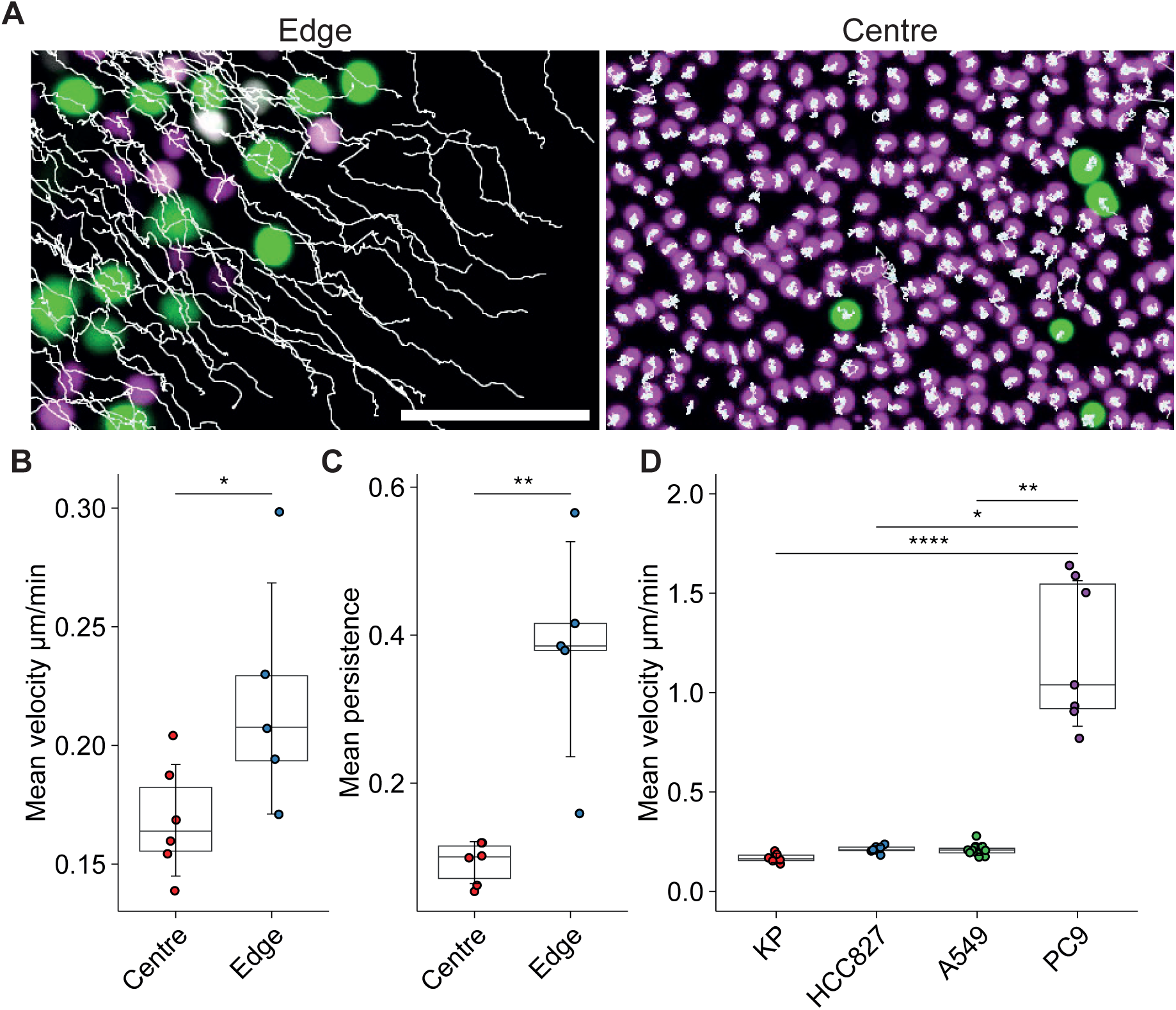
Analysis of migratory dynamics and forces. A. Representative images of central and edge regions from a KP-FUCCI single cell colony with cell tracks overlaid. Scale bar = 100 µm. B. Mean velocity of cells in the centre or edge regions where nuclei were tracked every 6 min for 18 h. C. Persistence of cells in the centre or edge regions where nuclei were tracked every 6 min for 18 h. Error bars are mean ± sd, pairwise comparisons by Wilcoxon signed rank test, * p < 0.05, ** p < 0.01, N = 4 independent experiments, n = 7 colonies. D. Quantification of mean cell velocity for cells in the centre of KP, HCC827, A549 or PC9 colonies where nuclei were tracked every 6 min for 18 h. Error bars are mean ± sd, pairwise comparisons by Dunn’s test with Benjamin-Hochberg p-value correction, * p < 0.05, ** p < 0.01, **** p < 0.0001, N = 3 independent experiments, KP n = 11, HCC827 n = 7, A549 n = 12, PC9 n = 12 colonies.

### Computational modelling links migratory dynamics to non-equivalent subclone fates

To explore the linkage between cell migration and subclone fates, we constructed a theoretical framework to explore the emergent relationships between these behaviours (Supplementary Methods). To this end, we generated an agent-based model, with each agent representing a cancer cell. Cells were capable of proliferation, migration, and interaction with neighbours. Migration was counteracted by an environmental drag term, reflecting the resistance of surrounding environment to cell movement. Neighbour interactions had both a short-range repulsive component to stop cells occupying the same space and a longer-range attractive component to reflect cell-cell adhesions (Figure 4a). Cells had a compressive stress threshold above which proliferation could not occur, reflecting the inverse correlation between density and proliferation that we observe. Model runs were initiated with a single cell and allowed to proceed until the colony size reached approximately 10^5^ cells. We varied combinations of intrinsic migratory potential, extrinsic drag, and cell doubling time (Figure 4). In the model runs with low cell migration, high extrinsic drag, and rapid proliferation, the colonies had an outer proliferative zone similar to that observed with KP cells (highlighted with blue boxes and lines in Figure 4b&c c.f. Figure 2a&b; Supp. Figure 4a&b). In addition, the centre of colonies showed elevated cell density (Figure 4d) and underwent a transition from initial exponential growth to a slower-than-exponential growth. This transition was linked to the emergence of a non-proliferative colony centre (Figure 4e, Supp. Figure 4h and Movie 4) and matched experimental analysis in Figure 2a&b. The emergent migratory dynamics of cells in the model also mirrored experiments with KP cells. Cells exhibited rapid linear outward migration at the colony edge and slow random migration in the colony interior (Figure 4f&g). Increasing internal cancer cell migration minimised differences in emergent migratory behaviour between the colony edge and interior (green boxes and lines Figure 4b-f). Reducing drag had a similar, but less pronounced, effect (yellow boxes and lines Figure 4f-g, Supp. Figure 4d and Movie 5), with greatly elevated proliferation in interior of tumour colony (Supp. Figure 4c). Thus, simply by tuning cell motility parameters we could recreate the diverse spatial patterns of proliferation observed experimentally (c.f. Figure 4b and 2h).

**Figure 4:**
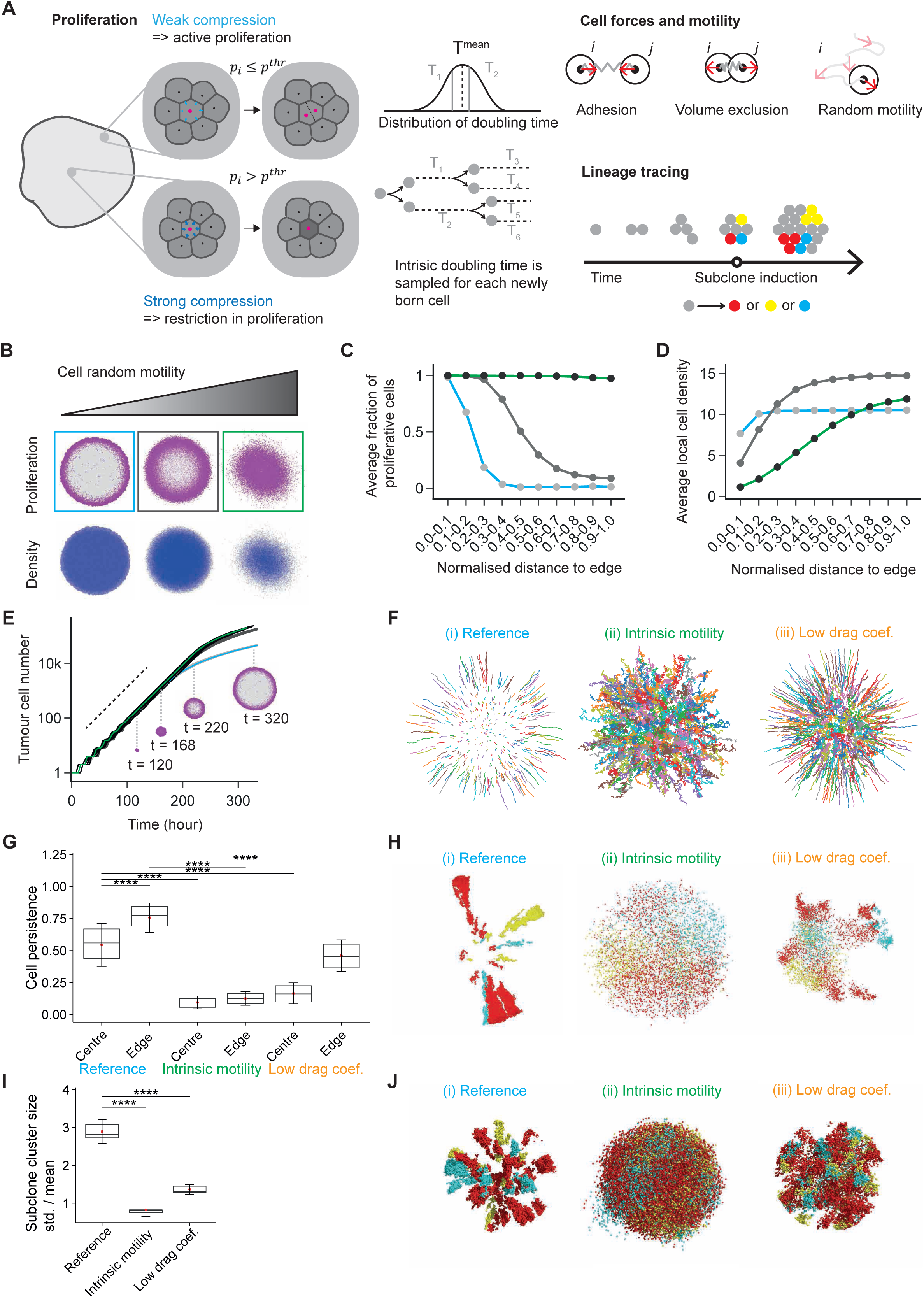
Computational modelling predicts cell migratory dynamics alters subclone evolutionary fates. A. Description of the agent-based model (ABM). The ABM characterises cell proliferation, adhesion and volume exclusion between cell neighbours, and cell random motility. A cell is assumed to become proliferatively arrested when it experiences strong compression by its neighbours. A lineage tracing protocol is implemented to label and track subclones in the model. B. Spatial patterns of proliferation and cell density in representative simulations with varying levels of random cell motility. Proliferatively active and arrested cells are labelled in magenta and light grey, respectively. Local cell density is labelled in blue, with greater intensity indicating higher density. Each column reflects snapshots from the same representative simulation. Boxes in different colours, which correspond to varying cell motility conditions and are consistently applied in the form of either boxes or curves in (C-E). Simulated colonies in these representative simulations have a size of about 4 mm. C. Average fraction of proliferative cells as a function of the distance to the tumour edge in simulations with varying model settings of cell random motility. In (C-D), the distance from a cell to the tumour edge is normalised to the largest distance in each simulation. Distances are then binned into 10 equally spaced radial zones to calculate intra-zone averages. Data points reflect the average of N = 16 simulations for each condition in (C-D). D. Average cell density as a function of the distance to the tumour edge in simulations with varying model settings of cell random motility. E. Tumour cell number over time, in simulations with different model settings of cell random motility. Solid lines reflect results from individual simulations, with N = 16 simulations for each motility condition. The dashed straight line is provided to indicate an exponential growth at early time points. Spatial patterns of proliferation from a single simulation at multiple time points are shown. F. Sampled cell trajectories over a duration of 18 hours in representative simulations with varying model settings – (i) “Reference” setting with low random cell motility and high drag coefficient; (ii) “Intrinsic motility” setting with high random cell motility and high drag coefficient; (iii) “Low drag coef.” setting with low random cell motility and low drag coefficient. These model settings are consistently applied in (F-J). More details about these model settings can be found in Supplementary Methods. G. Cell persistence over a duration of 18 hours at the colony centre versus edge, in simulations with different model settings of cell random motility and drag coefficient, respectively. N = 16 simulations for each condition. Welch two-sample t-test is performed. “****” reflects p <0.0001. For comparisons from left to right, t-values and degrees of freedom, (t, df), are: (-53.543, 4598.3), (130.56, 3086), (104.09, 3719.7), (254.61, 3771.5), (92.454, 5444.7). H. Spatial patterns of subclones in representative simulations with different model settings of cell random motility and drag coefficient. Subclone identities are labelled in red, yellow, or cyan. I. The standard deviation of subclone cluster size divided by the mean, compared between different model settings of cell random motility and drag coefficient. The subclone cluster size is measured as the number of cells in the subclone cluster divided by the total number of cells in the colony. In this analysis, a subclone is defined as a spatially contiguous patch of cells with an identical colour label (see Supplementary Methods). N = 16 simulations for each condition. Welch two-sample t-test is performed. “****” reflects p <0.0001. For comparisons from left to right, t-values and degrees of freedom, (t, df), are: (23.048, 23.723), (18.142, 19.843). J. Spatial patterns of subclones in three-dimensional realisations of representative simulations with different model settings of cell random motility and drag coefficient. Simulated colonies in these representative simulations have a size of about 2 mm. In (G) and (I), two ends of the box reflect the lower and upper quartiles, respectively, and the horizontal line dividing the box reflects the median. The red dot and error bar indicate the mean and the standard deviation, respectively.

To determine if the model was sufficient to explain the subclonal dynamics observed in our labelling experiments, we implemented *in silico* subclonal labelling. When the colony had reached a size of a few hundred cells a small percentage of cells were irreversibly assigned a colour label that would be inherited by their progeny. This enabled the fate of these *in silico* subclones to be tracked. Figure 4h shows the visual outputs of these simulations, revealing a high level of concordance with our experimental observations of KP cells when cell motility was low and drag was high (henceforth we refer to this model condition referred to as “Reference”). This concordance was also evident in the relationship between subclone size and position relative to the boundary (quantified in Figure 4i and Supp. Figure 4h). Similar conclusions were obtained if subclones were determined simply based on contiguous patches of colour, as is done experimentally, or on true lineage, which could be tracked in the model (compare Figure 4i and Supp. Figure 4e). Thus, our analyses are not confounded by subclone fission and fusion events.

We next investigated the predicted patterns of subclone labelling in simulations with high cell motility (referred to as “Intrinsic motility” with green text) and lower drag (referred to as “Low drag coef.” With yellow text). These revealed that the ‘streaks and spots’ pattern was disrupted when cell migration was elevated. Instead subclones appeared more diffuse and without clear boundaries between subclones; in other words, high motility or low drag led to greater subclone mixing (quantitatively demonstrated in Supp. Figure 4f-g using a metric of subclone overlap). More significantly, the size distribution of subclones in conditions of high migration and mixing was much more homogenous than in low migration conditions (Figure 4i and Supp. Figure 4e). Adaptation of our model to three spatial dimensions further confirmed its ability to replicate the subclonal patterns observed in KP spheroid grown in agar, with high cell migration disrupting this pattern (Figure 4j c.f. Supp. Figure 1d). Together, these data demonstrate that the patterns of clonal growth that we observe in different lung cancer models can be explained simply by the interplay of cell proliferation, cell migration dynamics, and compressive stress suppressing cell proliferation (Supp Figure 4i). Moreover, the inequality in subclone fates observed in the KP model is predicted to be reduced if migration is increased.

### Enhancing cell migration to equalises subclonal fates

We next tested if the variable subclone fates in the KP model were reduced by increasing cell migration. We employed two strategies to increase the migration of KP cells: TGFβ treatment and Rab5A over-expression. This choice was motivated by the ability of TGFβ to drive epithelial cancer cells to a migratory and mesenchymal phenotype (Oft *et al*., 1996; Oft, Heider and Beug, 1998) and the ability of Rab5A to counter-act the phenomenon of cell jamming (Malinverno *et al*., 2017; Palamidessi *et al*., 2019), which is the cessation of cell migration at high cell density. Figure 5a&b show that TGFβ treatment does indeed promote cell migration and the loss of cell-cell adhesions, which is analogous to increasing the cell motility term in our computational model. Moreover, TGFβ treatment also increased the frequency of cell migration tracks crossing, which reflects cells changing position relative to one another and mixing within the colony (Figure 5c&d). Increased cell migration is predicted to reduce the proliferative arrest in the centre of the colony and reduce the variability in subclone fate (Figure 4b-i). Strikingly, TGFβ increased EdU incorporation in the colony centre and disrupted the spots and streaks pattern of subclones normally observed in our KP lung colonies (Figure 5e). This was reflected in a pronounced reduction in the standard deviation of subclone size (Figure 5f). To confirm that the effect of increasing migration on subclonal dynamics was not specific to TGFβ treatment, we also analysed the effect of Rab5A over-expression, which can drive the migration of epithelial cells (Malinverno *et al*., 2017; Palamidessi *et al*., 2019). This led to altered subclonal patterns, with a reduced standard deviation in subclonal patch sizes observed in Rab5A over-expressing tumour colonies (Supp. Figure 5a-c). This change was also associated with altered shapes subclonal patches, reminiscent of those observed in low drag coefficient simulations, and increased proliferation in the colony centre. Together, these data demonstrate that cancer cell migration suppresses the variability in the fate of subclones with neutral changes.

**Figure 5:**
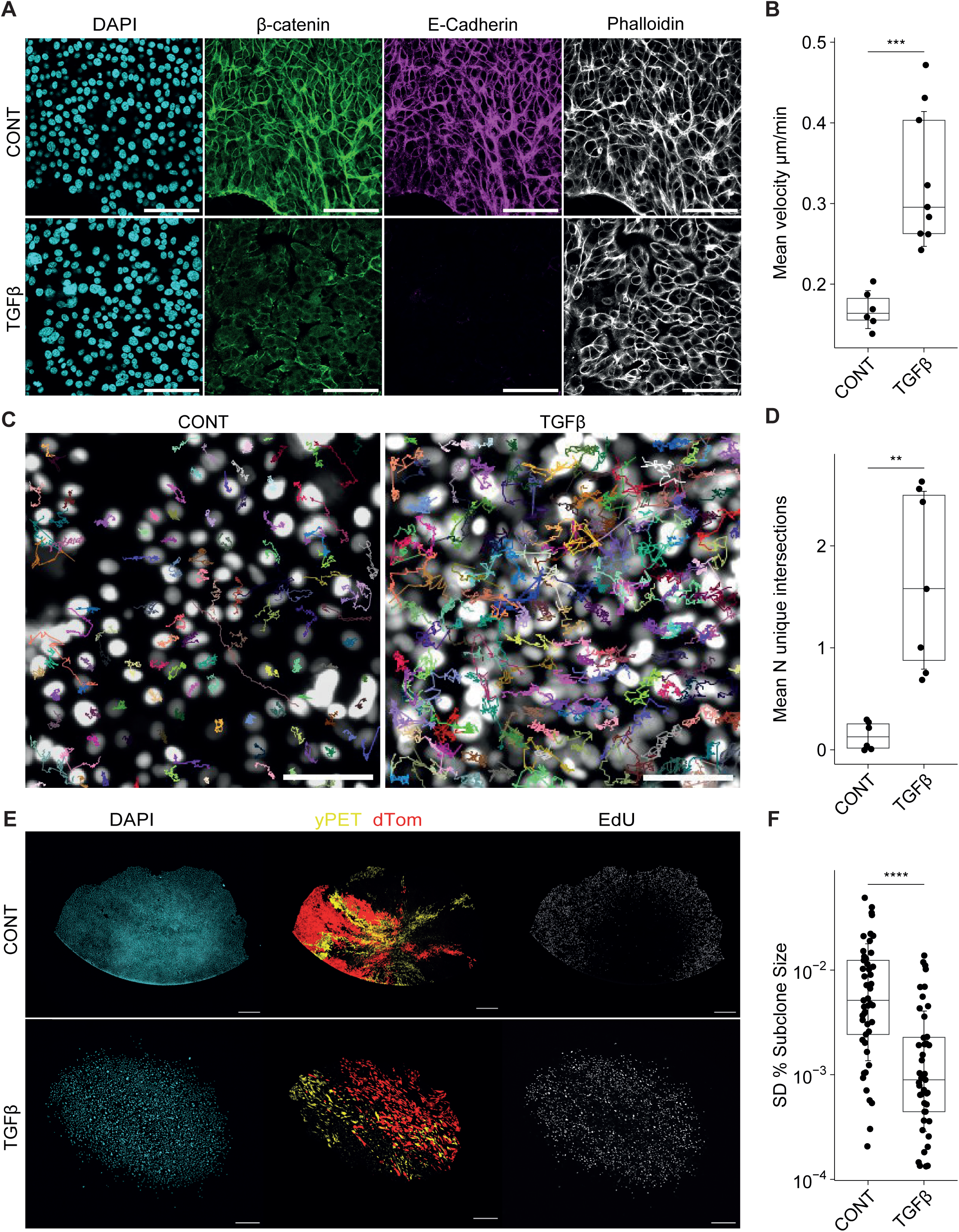
Cell migration alters subclonal fates. A. Representative immunofluorescence staining of DAPI, β-catenin, E-Cadherin and Phalloidin in KP control and TGFβ treated single cell colonies at the boundary. Scale bar = 100 μm. B. Quantification of mean velocity for cells in the centre of KP control and TGFβ treated colonies where nuclei were tracked every 6 min for 18 h using StarDist-TrackMate. Error bars are mean ± sd, pairwise comparisons by Wilcoxon signed rank test, *** p < 0.001, N = 4 independent experiments, control n = 6, TGFβ n = 9 colonies. C. Representative images of central regions from KP control and TGFβ treated colonies with tracks overlayed. Scale bar = 100 µm. D. Quantification of cell mixing using number of unique track intersections in central regions from KP control and TGFβ treated colonies. First, a convex hull was constructed for each track to define a cell domain then the number of different tracks crossing into that area was quantified to give number of unique intersections. N = 3 independent experiments, control n = 6, TGFβ n = 6 colonies. Error bars are mean ± sd, pairwise comparisons by Wilcoxon signed rank test, ** p < 0.01. E. Representative images of KP-pOncobow control and TGFβ treated colonies. Single KP-pOncobow cells were plated and treated with 2 nM 4-OHT for 6 h on day 3. 1 ng ml^-1^ TGFβ or DMSO control were added on day 5. Cells were then left in culture until day 14 when a 1 h 10 μM EdU pulse was performed prior to fixation and immunostaining for DAPI and EdU. Scale bar = 500 μm, colours represent DAPI (cyan), yPET (yellow) and dTomato (red) subclones, EdU (grey). F. Quantification of standard deviation of subclone size normalised to colony area. Error bars are mean ± sd, pairwise comparisons by Wilcoxon signed rank test, **** p < 0.0001, control n = 43, TGFβ n = 49 colonies, N = 8 independent experiments.

### TGFβ and EMT are linked to subclonal mixing in human lung cancer

Thus far, we have focused on experimental models of lung cancer. We next sought to determine what the level of mixing might be in human lung adenocarcinoma. While it is not possible to perform subclonal labelling in cancer patients, metrics such as the distribution of variant alleles frequencies (VAF) contain information about the subclonal composition of tumours (Ding *et al*., 2012; Gerlinger *et al*., 2012; Klco *et al*., 2014; Jamal-Hanjani *et al*., 2017; McGranahan and Swanton, 2017). To relate observations made in our models, both experimental and *in silico*, we implemented a mutational process in our computational model and developed a computational framework to infer subclone mixing based on features of variant alleles (Supplementary Methods). For simplicity, each cell was considered to have 10 000 alleles and these could become mutated in a manner linked either to cell proliferation or simply the passage of model time. Thus, cells in the model would accumulate mutant alleles over time. Instead of analysing the pattern of coloured clones, we analysed the frequency distribution of mutant alleles in simulated regional biopsies - analogous to the sampling process used in the TRACERx lung cancer study (Jamal-Hanjani *et al*., 2017; Frankell *et al*., 2023) (Figure 6a). Increasing internal cell motility or reducing external drag, both of which promote subclone mixing (Supp. Figure 4f-g), generated different and characteristic variant (mutant) allele frequency plots. In particular, low cell migration and consequently high variability in the fate of neutral subclones resulted in a larger number of ‘mutant alleles’ undergoing expansion with VAF values between 0.1 -0.4 (given the acronym MUE for Mutations Under Expansion – Figure 6b). The difference between non-mixed (“Reference”) and mixed tumour models (“Intrinsic motility” or “Low drag coef.”) was apparent regardless of whether mutations were coupled to cell proliferation or simply accumulated with time (Supp. Figure 6a-b). Furthermore, the differences were also maintained if some of the mutations carried either a selective advantage or disadvantage (Supp. Figure 6a-b).

**Figure 6:**
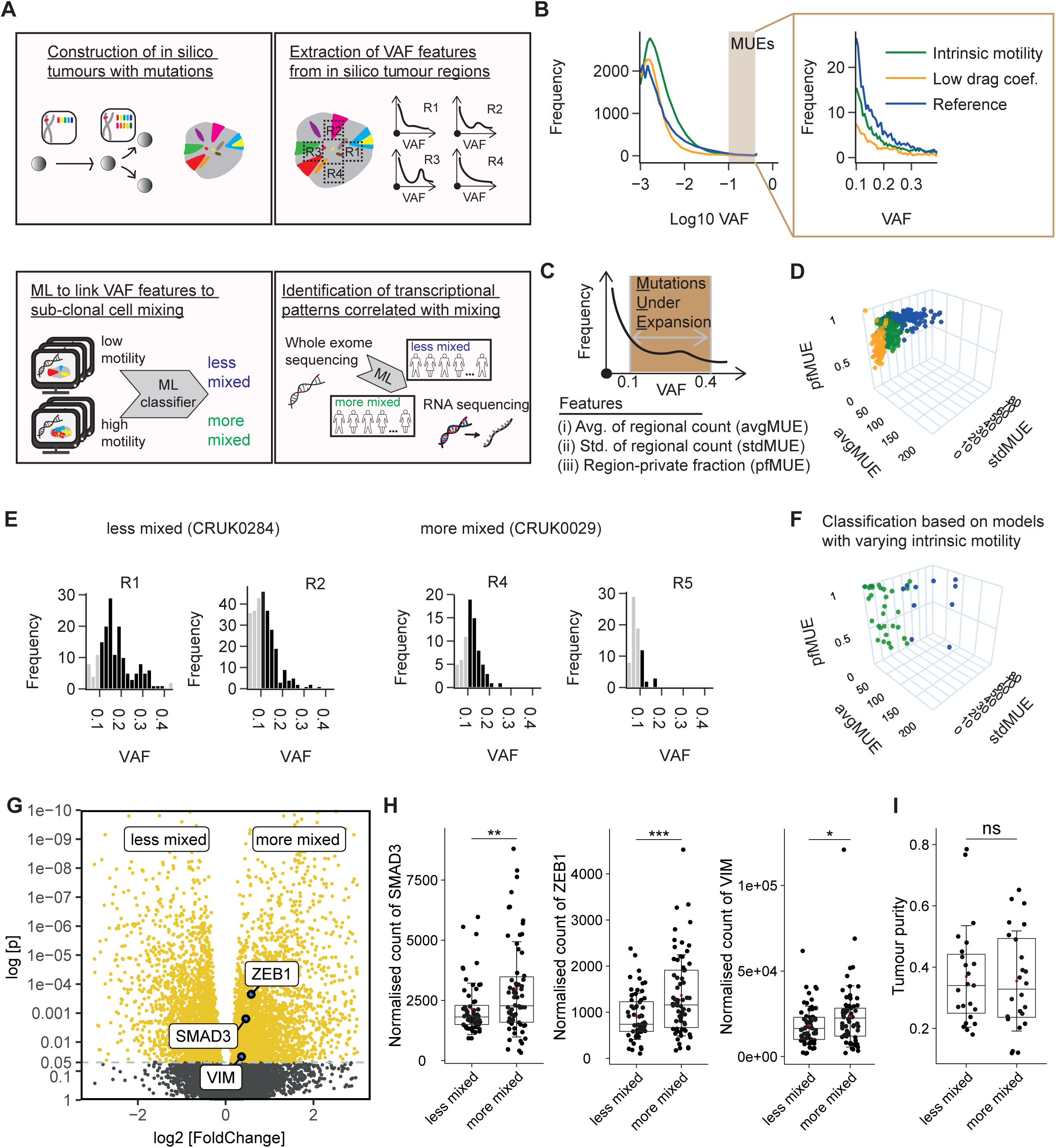
Computational modelling links TGFβ and EMT with subclonal mixing in human lung cancer. A. Schematic illustrating the steps to infer the degree of subclonal mixing in human lung cancer. More details can be found in Supplementary Methods. B. Frequency distributions of regional variant allele frequency (VAF) in simulations with varying model settings of random cell motility and drag coefficient, labelled as “Reference” (blue; high drag coef. And low motility), “Intrinsic motility” (green; high motility), and “Low drag coef.” (orange; low drag coef.), respectively. Lines reflect frequency distribution of VAFs in simulated tumours, averaged across regions of N = 120 simulations per condition. The distribution of Log10-transformed VAF over a wider range is shown in the left panel; the distribution of VAF between [0.1, 0.4], which defines Mutations Under Expansion (MUEs), is shown in the right panel. C. A schematic illustrating extraction of quantitative features from VAF distribution. These features are based on MUEs, which reflect mutant alleles with a VAF between 0.1 and 0.4 in this analysis. D. VAF features of simulated tumours with different model settings of random cell motility and drag coefficient. N = 120 simulations for each condition. E. Frequency distributions of VAF in two exemplar regions of two LUAD tumours inferred to be “less mixed” and “more mixed”, respectively. VAF distribution in two regions is shown for each tumour. Bars that reflect VAFs within the interval of 0.1 and 0.4 are in black. F. Inferred mixedness of LUAD tumours with predominantly solid histological subtype in the TRACERx Lung study plotted in the VAF feature space, based on a Support Vector Machine (SVM) classifier trained using simulations with low vs high levels of random cell motility. Tumours with more extreme MUE features are not presented in this plot (Refer to Supp Figure 6c and Supplementary Table 1). G. Differential expression of genes between subsets of tumours inferred to be “more mixed” (N = 66 regions from 20 tumours) and “less mixed” (N = 60 regions from 20 tumours). Selected significantly differentially expressed epithelial-to-mesenchymal transition (EMT) genes are highlighted in the plot. H. Normalised expression of EMT marker genes in subsets of tumours inferred to be “less mixed” and “more mixed”, respectively. Welch two-sample t-test is performed. “***” reflects p<0.001; “**” reflects p<0.01; “*” reflects p<0.05. For comparisons from left to right, t-values and degrees of freedom, (t, df), are: (-2.8963, 101.77), (-3.5913, 105.32), (-1.9834, 110.78). I. Tumour purity in subsets of tumours inferred to be “less mixed” (N = 24 tumours) and “more mixed” (N = 23 tumours), respectively. Welch two-sample t-test is performed. “ns” reflects not significant, p = 0.7625. The t-value and degree of freedom, (t, df), is: (0.30405, 44.849). In (H-I), two ends of the box reflect the lower and upper quartiles, respectively, and the horizontal line dividing the box reflects the median. The red dot and error bar indicate the mean and the standard deviation, respectively.

We utilised the information from our model about the relationship between cell migration and variant-allele frequencies to determine levels of mixing in lung adenocarcinoma from the TRACERx study, which incorporates multi-region sampling of the tumour genome (Jamal-Hanjani *et al*., 2017; Frankell *et al*., 2023). VAF distributions from poorly- and well-mixed simulations could reliably be distinguished by the combination of three metrics – mean number of regional MUEs, standard deviation of MUEs, and fraction of region specific MUEs (Figure 6c&d). These three metrics derived from *in silico* simulations with known levels of mixing, were then used within a support vector machine to classify VAF distributions in 40 lung adenocarcinoma with solid histology in the TRACERx cohort into those with similar characteristics to poorly- and well-mixed simulations. Figure 6e&f show the similarity in MUE features between our simulations and the clinical data and that we can infer differences in mixing between human lung adenocarcinoma. Tumours with high or low inferred mixedness had similar overall tumour purity (Figure 6i, Supp Figure 6g, Supplementary Table 1). To explore factors that might drive differing degrees of cell mixing, we compared both the mutational profiles and the transcriptomes of lung tumours with the different classes of VAF distribution. No clear associations between driver mutations and inferred mixing were observed (Supplementary Table 2). Notably, increased EMT and TGFβ-associated gene expression was observed in tumours inferred to have high levels of cell mixing (Figure 6g-h). Concordant results were obtained if inference of subclone mixing and transcriptomic analysis were based on simulations with reduced drag coefficient (Supplementary Table 1, Supp Figure 6d-g). These data demonstrate the relevance of findings from our model regarding TGFβ-driven EMT suppressing the variability in subclone fate in human tumours.

### Cancer-associated fibroblasts modulate subclonal dynamics

The tumour microenvironment modulates many features of cancer cell behaviour, including cell migration and invasion (Olumi *et al*., 2000; Gaggioli *et al*., 2007; Karnoub *et al*., 2007; Kuzet and Gaggioli, 2016). We therefore asked if there might be links between stromal cell types and subclonal dynamics. Intriguingly, we observed increased expression of several genes indicative of cancer-associated fibroblasts, including ACTA2 (αSMA), PDGFRB, THY1, and COL1A2, in tumours predicted to have high levels of mixing based on their VAF characteristics (Figure 7a-b, Supp. Figure 7a-b). Overall tumour purity was equivalent between tumours with high and low predicted mixing, thus the increased expression of CAF markers is not an artefact of low tumour purity (Figure 6i, Supp Figure 6g). Given the extensive literature on cancer-associated fibroblasts promoting cancer cell migration and invasion, we tested whether stromal fibroblasts could modulate subclonal dynamics by directly adding human lung cancer-associated fibroblasts into our tumour colony assays. This led to alterations in tumour colony shape and increased cancer cell migration (Supp. Figure 7c&e, Figure 7c&d) Moreover, cancer-associated fibroblast reduced the variability between subclone sizes (Figure 7e), providing experimental corroboration of the inference from patient data that stromal fibroblasts influence subclonal dynamics.

**Figure 7:**
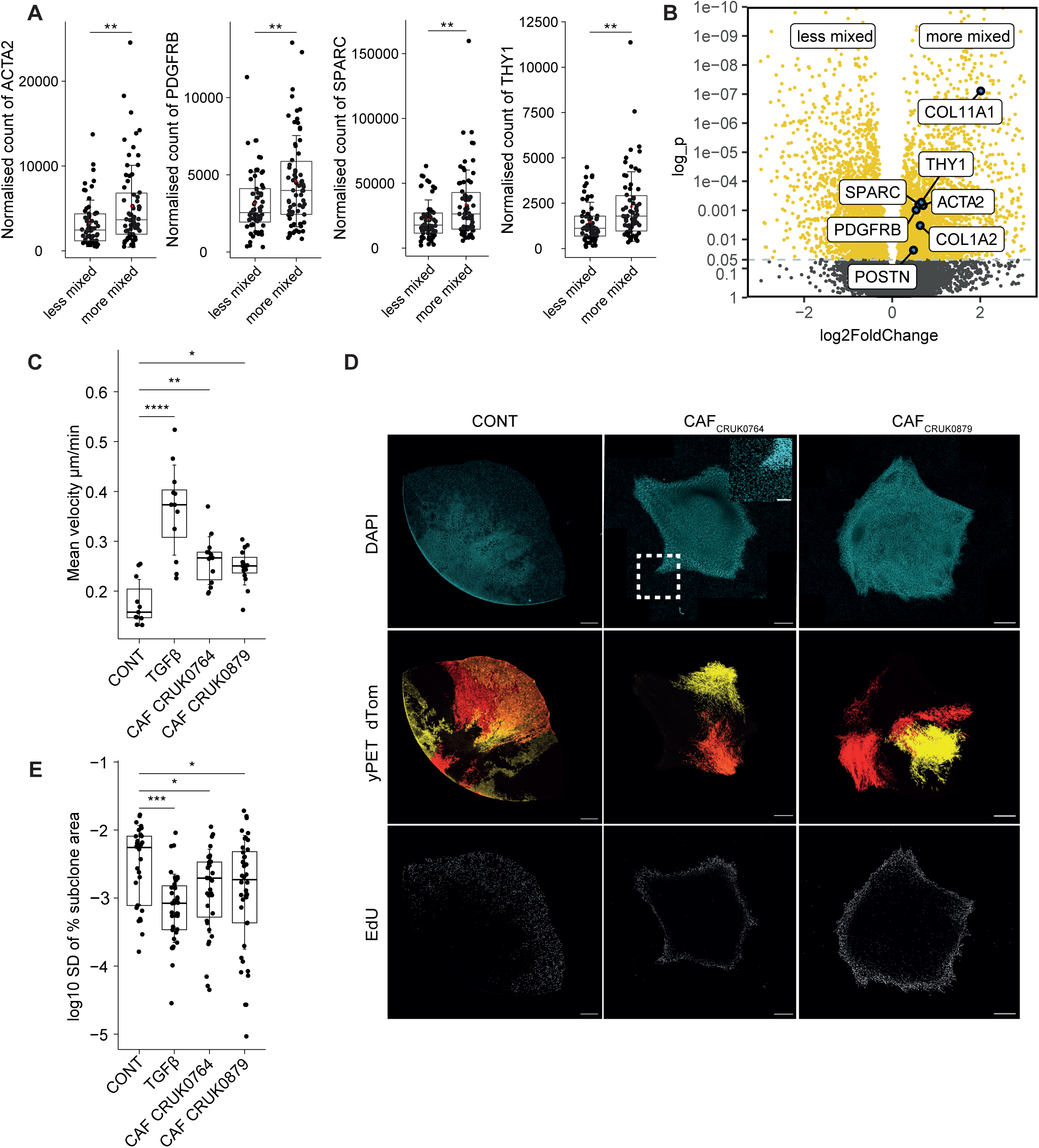
Cancer associated fibroblasts modulate subclonal mixing. A. Normalised expression of CAF marker genes in subsets of tumours inferred to be “less mixed” and “more mixed”, respectively. Inference of mixedness was based on a SVM classifier trained using simulations with low vs high levels of random cell motility. B. Differential expression of genes between subsets of tumours inferred to be “more mixed” (N = 66 regions from 20 tumours) and “less mixed” (N = 60 regions from 20 tumours). Select significantly differentially expressed CAF and ECM marker genes are highlighted in the plot. C. Quantification of velocity in ROIs at the colony centre and edge of KP-pOncobow control, TFGβ, KP-CRUK0764 and KP-CRUK0879 colonies on day 11 of growth. Cell migration was tracked using StarDist-TrackMate for 108 frames with 10 min intervals, control n= 11, TGFβ n = 12, KP-CRUK0764 n = 14, KP-CRUK0879 n = 15 colonies per condition, N = 4 independent experiments. D. Representative confocal images of colony subclone patterns in control, KP-CRUK0764 and KP-CRUK0879 co-culture, colours represent DAPI (cyan), yPET (yellow) and dTomato (red) subclones, EdU (grey). Scale bar = 500 μm, E. Quantification of standard deviation of subclone size normalised to colony area. Control n = 29, TGFβ n = 39, KP-CRUK0764 n = 35, KP-CRUK0879 n = 40 colonies per condition, N = 4 independent experiments. Error bars are mean ± sd, pairwise comparisons by Dunn’s test with Benjamin-Hochberg p-value correction, * p < 0.05, ** p < 0.01, *** p < 0.001, **** p < 0.0001.

### Changing migratory dynamics alters the rate at which aggressive subclones become dominant

Finally, we sought to explore how inter-cellular mixing might affect the behaviour of subclones with selective advantage or disadvantage. We returned to our *in silico* model to interrogate the effect of altering either the growth rate or therapy responsiveness of subclones. To model growth advantage, 5 out of the 10 000 alleles, when mutated, lead to faster proliferation (with the average doubling time of cells harbouring these mutant alleles reduced by 20%). The model was then run with either high or minimal levels of cell mixing (resulting from increased intrinsic cell motility), with the proportion of cells with mutant alleles conferring shorter cell cycle time tracked (Supp. Figure 8a&b). Supp. Figure 8c shows that tumours with the largest sweep of advantageous subclones had low levels of cell mixing. The benefit arose in simulations where the advantageous subclones emerged at early timepoints prior to the onset of the density induced inhibition of proliferation (time of subclone birth is colour coded in Supp. Figure 8c). These data argue that the unequal fate of subclones in the absence of cell mixing permits advantageous mutations that arise early and in a favourable location to dominate more rapidly. Consistent with this, the expression of genes correlated with low levels of cell mixing were associated with worse overall survival in patients with stage I disease, but not higher stage disease (Supp. Figure 8d, Supplementary Table 3)

Therapy resistance was modelled by eliminating a fixed proportion of cells at each time step, with 5/10 000 alleles, when mutated, conferring resistance to elimination. Simulations were initiated without cell elimination and it was triggered at a certain size threshold – mimicking the initiation of treatment in an established tumour (Figure 8a). Once again, high and low levels of cell mixing were explored. In contrast to the growth advantage simulations, high levels of cell mixing consistently favoured the sweep of subclones resistant to cell killing (Figure 8b&c). In conditions of minimal migration, subclones resistant to killing still became dominant, but at a slower rate and with sensitive streaks of cells persisting. This transition to dominance was also associated with changes in the colony morphology, with expanding resistant subclones forming bulges at the colony edge (Figure 8b, Movie 6). These data indicate that the relationship between cell mixing, subclonal dynamics, and the rate of clonal sweeping differs depending on whether cells are being selected for growth advantage or resistance to cell death.

**Figure 8:**
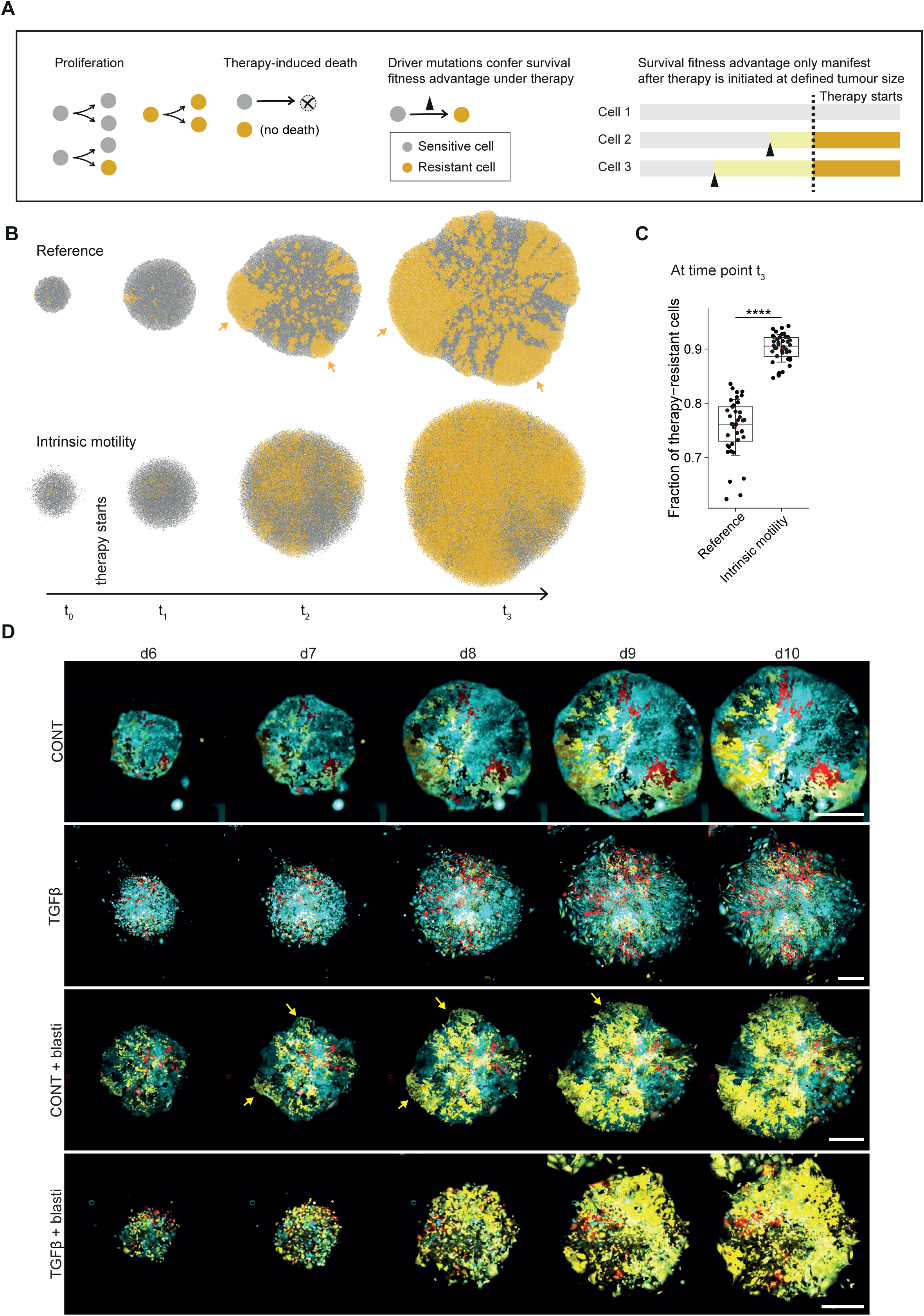
Computational modelling links subclonal mixing with differential outgrowth rates of fitter subclones. A. Schematics illustrating model implementation of survival fitness advantage. Therapy-resistant cells (yellow) emerge from acquiring a driver mutation upon proliferation. Survival fitness advantage only manifest in cells harbouring the driver mutation after therapy is initiated, when the tumour reaches N=10000 cells in the model. 5 out of 10000 alleles, when mutated, confer a survival advantage under therapy. Mutations of these alleles are referred to as driver mutations. B. Spatial patterns of therapy-resistant cells (yellow) over time in representative simulated tumours with low and high random cell motility, respectively. C. Fraction of therapy-resistant cells at a late stage of simulated tumours with low and high random cell motility, respectively. N = 40 simulations for each condition. Welch two-sample t-test is performed. “****” reflects p <0.0001. The t-value and degree of freedom, (t, df), is: (-16.039, 57). D. Representative time lapse images of KP-pOncobow-BSR colonies every 24 h from day 6 to day 10 of growth. Colours represent subclones H2B-mTurquoise (cyan), yPET-BSR (yellow), dTomato (red). Briefly, single cell colonies were plated, recombination induced on day 3 with 6 h 2 nM 4-OHT, 1 ng ml^-1^ TGFβ was added on day 5 then 5 µg ml^-1^ Blasticidin (blasti) was added on day 6 at least 6 h prior to imaging. Scale bar = 500 μm, arrows denote bulging subclones. N = 3 independent experiments, control n = 47, control + blasti n = 53, TGFβ n = 58, TGFβ + blasti n = 45 colonies per condition. In (C), two ends of the box reflect the lower and upper quartiles, respectively, and the horizontal line dividing the box reflects the median. The red dot and error bar indicate the mean and the standard deviation, respectively.

To explore the model prediction regarding subclone migration and the rate of therapy resistant subclone dominance, we selected blasticidin and the bsr gene (Blasticidin S-resistance). This choice was motivated by resistance being conferred by a single gene and our desire to avoid the well-documented and complex interaction of TGFβ, EMT, and conventional chemotherapy (Yang *et al*., 2006; Cheng *et al*., 2007; Kudo-Saito *et al*., 2009; Thiery *et al*., 2009; Debaugnies *et al*., 2023). We modified our Oncobow cassette to couple the expression of yPET and a blasticidin resistance gene. We then repeated our colony assays in the presence of combinations of TGFβ to drive subclone migration and blasticidin to apply therapeutic selection pressure. Figure 8d & Movie 7 show that yPET expressing subclones were neither favoured or disadvantaged in the absence of blasticidin. In contrast, when blasticidin was added the yPET expressing subclones became dominant. In the absence of TGFβ, expanding yPET subclones formed bulges at the colony edge and some sensitive cells remained in streaks – thereby confirming predictions of the agent-based model. These data demonstrate that TGFβ-driven neutralisation of the ‘place of birth’ effect favours therapy resistance.

## Discussion

Evolutionary principles underlie both the development of cancer and the emergence of acquired resistance to therapy (Greaves and Maley, 2012; Lloyd *et al*., 2016; Sun *et al*., 2017; Zahir *et al*., 2020). Much effort has been devoted to understanding the mutational processes giving rise to genetic variation and which mutations confer advantages to cancer cells (Gerlinger *et al*., 2012; Nik-Zainal *et al*., 2012; De Bruin *et al*., 2014; Williams *et al*., 2016, 2018; Jamal-Hanjani *et al*., 2017; Gerstung *et al*., 2020). The advantage or disadvantage of a given mutation is also dependent on the environmental context, such as the extent of immune surveillance (Janiszewska *et al*., 2019; Lakatos *et al*., 2020; Dhainaut *et al*., 2022; Risom *et al*., 2022) or presence of drug (Hata *et al*., 2016; Gallaher *et al*., 2018). In other areas of evolutionary analysis, the number of individuals in the population, their migration, and mixing are also well-established to influence evolutionary dynamics (Schreck *et al*., no date; Fusco *et al*., 2016; Farrell *et al*., 2017; Kayser *et al*., 2018; Paulose and Hallatschek, 2020). These factors have also been considered in models of cancer evolution (Waclaw *et al*., 2015; West *et al*., 2021), but demonstration of their role in experimental and clinical contexts is scarce. In this work, we employ integrated experimental, computational, and clinical analyses of lung adenocarcinoma to discover that in the absence of cancer cell migration, the long-term fate of subclones is heavily influenced by their place of birth. Notably, gene expression linked to low levels of subclone mixing was associated with worse outcomes in early stage disease (Supp. Figure 8d). This suggests that ‘born to be bad’ tumours arise from a combination of early unfavourable genetic events (Sottoriva *et al*., 2015) and the location and tissue dynamics in which those genetic events occur. As tumours develop and begin to be treated, the relationship between cell mixing and bad outcomes switches. In tumours subject to cytotoxic therapies, the migration and mixing of cancer cells, possibly resulting from either cancer cell EMT or the presence of stromal fibroblasts, favours the dominance of intrinsically therapy resistant subclones.

Lineage tracing of subclone fate was central to our experimental analysis. This approach has been extensively applied to tissues with rapidly proliferating stem cells and high cell turnover; such as the skin and intestine (Lamprecht *et al*., 2017; Lenos *et al*., 2018; Reeves *et al*., 2018; Van Der Heijden *et al*., 2019; Flanagan *et al*., 2021). These analyses have demonstrated that gain of an oncogenic mutation in a stem cell is much more likely to become fixed and generate a tumour. Factors that enhance stem cells, such as SPP1 produced by fibroblasts, also favour tumorigenesis and the expansion of subclones (Lenos *et al*., 2018). These analyses have also provided support for low levels of subclone mixing in early/small squamous cell carcinoma, with an increase in more advanced lesions (Reeves *et al*., 2018). However, the situation in tissues that are largely quiescent during homeostasis is much less clear. The alveolar regions of the lung are one such quiescent tissue. If damage occurs, then regeneration results from the re-entry of alveolar type II (ATII) cells into the cell cycle and, as a result, these cells are considered to be stem cells in lung (Bender Kim *et al*., 2005). Consistent with this, and with analysis of the skin and intestine, ATII cells are thought to be the cell of origin of LUAD (Thaete and Malkinson, 1991; Jackson *et al*., 2001). We propose that the proliferation of LUAD cells is heavily influenced by their density and the compressive stress that they experience. The mechanism by which compressive stress reduces proliferation remains to be determined. It may relate to the thermodynamic challenge of moving the osmolytes required for growth against a pressure gradient, which has been demonstrated in yeast (Alric *et al*., 2022). We think it is unlikely to involve canonical cell cycle controls, such as Rb and p53, as these are lost or de-regulated during the transformation process and all LUAD cell lines tested exhibit reduced proliferation at high density (Delarue *et al*., 2014; Streichan *et al*., 2014; Zatulovskiy *et al*., 2020; Devany *et al*., 2022). It is also distinct from contact inhibition as cells surrounded on all sides continued to proliferate if their density remained low. When considered alongside cell migration, the phenomenon of density induced proliferation restriction leads to emergent patterns of proliferation and subclonal dynamics. When migration is low, tumours develop non-proliferative centres and only subclones originating near the edge can become established. This is consistent with the observation of compressive stress in the interior of tumours in pre-clinical studies (Delarue *et al*., 2014; Streichan *et al*., 2014; Di Meglio *et al*., 2022), and with the observation of lower Ki-67 expression at the tumour core in comparison to periphery revealed by digital spatial profiling of LUAD samples (Tavernari *et al*., 2021). If migration is high, this provides a means to dissipate the compressive stress and lower cell densities leading to more uniform distribution of proliferating cells and subclone fates. Stromal fibroblasts also promote the migration of cancer cells and more equivalent subclone fates. However, the spatial patterns of subclones following TGFβ treatment and CAF co-culture were different (compare Figures 5&7), suggesting that CAFs are not acting simply as a source of TGFβ.

The structure and organisation of human lung cancer is complex; nonetheless, we propose that the same factors act to govern where cancer cells can proliferate. This is supported by the concordance between our model systems and human patient data (Figure 7). The structural and mechanical properties of the lung tissue surrounding human tumours is more varied and complex than in our models. This, coupled with slower proliferation rates in human disease means that patterns arising in human disease are more complex than in our models. Nonetheless, increased proliferation at the periphery of tumours is observed in many tumours. In our 2D in vitro analyses, all cells have similar access to nutrients and oxygen in the media. However, in vivo these factors can become limiting in the centre of tumours with central necrosis resulting. We predict that cell death in the tumour centre would relieve the compressive stress and lead to a resumption of proliferation – conceptually similar to the experiments in which we removed a central patch of the tumour colony. This is supported by recent analysis of clear cell renal cell carcinoma (ccRCC) that observed a subset of tumours with peripheral proliferative zones and quiescent centres, and other tumours with central necrosis and increase cell turnover (Zhao *et al*., 2021).

To summarise, through combined experimentation, computational modelling, and clinical analysis, we demonstrate that EMT and cell migration promoted by stromal fibroblasts fundamentally alters the competitive dynamics between subclones in tumours. This has important consequences for the rate at which therapy resistant subclone become dominant. Moreover, our framework relating migratory dynamics, the build-up of compressive stress, subclonal fates and competition has the power to explain multiple facets of cancer evolution.

## Methods

### DNA constructs

The pOncobow constructs are modified versions of the pMAGIC cytobow that was a gift from Dr. Karine Loulier (INSERM, France). To generate pOncobow *(ITR-IoxP-loxN-lox2272-H2B-mTurquoise2-IRES-PURO-STOP-loxP-dTomato-FLAG-STOP-loxN-mKeima-V5-STOP-lox2272-yPET-HA-STOP-ITR)* from pMAGIC cytobow *(ITR-IoxP-loxN-lox2272-H2B-eBFP-STOP-loxP-dTomato-STOP-loxN-mCerulean-STOP-lox2272-yPET-STOP-ITR),* single colour constructs, tDTomato (Shaner *et al*., 2004), mKeima (Kogure *et al*., 2006), and yPET (Nguyen and Daugherty, 2005) were flanked by a flexible linker sequence (3x[SGGGG]) and biochemical epitopes FLAG, V5 or HA respectively. In addition, restriction sites were added flanking each fluorescent protein spaced 10-15bp apart from each other and the start codon of a fluorescent protein. A puromycin selection cassette was placed 3’ to H2B-mTurquoise2 with an intervening internal ribosome entry sequence (IRES) sequence to decouple continuous translation of H2B-mTurquoise2 and puromycin expression. Due to the highly repetitive nucleotide sequences flanking fluorescent proteins and LoxP/N/2272 sequences in the pMAGIC construct both an iterative Gibson cloning process, and a nucleotide sequence optimization process for inserted vectors was required to ensure that the greatest number of single cutting restriction sites would be made available for the final construct. The sequences of dsDNA geneblocks vectors (IDT, Custom Order) and primers (Sigma, Custom Order) used to create pOncobow5 can be found in table 1. A typical cloning process consisted of a plasmid digest using restriction enzymes (all enzymes were purchased from NEB, using high-fidelity (HF) enzymes where available) followed by agarose gel electrophoresis, gel extraction, and subsequent gel purification (Thermo-Fisher, Cat. #K0691) to isolate cut plasmid DNA. Insert was PCR amplified from other plasmids or was purchased as dsDNA (IDT, See Table). Following gel purification of PCR amplified DNA or preparation of dsDNA geneblocks as per manufacturer’s instructions (IDT), cut plasmid DNA and PCR or dsDNA insert were recombined using Gibson cloning (NEB, Cat. #E5510), relying on a homology of 25bp between flanking ends of cut plasmid DNA and PCR or dsDNA to facilitate the homologous recombination process.

### Cell line generation, validation, and maintenance

The KPCreERT^2^ (KP) cell line used in this study was a gift from the Downward Lab, Francis Crick Institute and was derived from a lung tumour induced by intranasal administration of adenoviral FLP-FRT recombinase in a female C57BL/6 CreERT^2^; Trp53^frt/frt^; KRas^FSF-G12D^ mouse as previously described (DuPage, Dooley and Jacks, 2009). Stable transfection of KP, PC9, A549 or HCC827 cells was performed by mixing plasmid DNA comprised a 1:1 mixture PiggyBac retrotransposase plasmid and a single plasmid of interest (pOncobow, single gene expressing constructs, pFastFUCCI (Koh *et al*., 2017), EKAREV (Komatsu *et al*., 2011)) with 15 µl Attractene (Qiagen, Cat. #301005) or 10 µl lipofectamine 2000 (Invitrogen, Cat. #11668019) and incubating the mixture for 30 minutes on ice. During this time, 5 x 10^5^ cells on 100 mm^2^ in tissue culture dishes or 2 x 10^5^ cells in 6-well plates were washed with PBS twice and replaced with OptiMEM (Gibco, Cat. #31985). Following mixture incubation, transfection mix was then added dropwise to the dish. Following 24 h of uninterrupted incubation, cells were then replaced with DMEM (Gibco, Cat. #10569010) with 20% Fetal Bovine Serum (FBS) (Gibco, Cat. #16000069) for KP cells, DMEM; 10% FBS for A549 cells or RPMI (Gibco, Cat. #21875091); 10% FBS for PC9 and HCC827 cells. Once cells appeared viable and fluorescent clones were visible, the media was changed to the appropriate culture media supplemented with 1% Penicillin and Streptomycin (P/S) (Gibco, Cat. #15140122). Cells were then selected in 2 µg ml^-1^ puromycin or for EKAREV cell lines 2 µg ml^-1^ Blasticidin and sorted via FACS to obtain a pure population of cells.

For lentiviral infections, 2 x 10^6^ 293T packaging cells were plated in a 100 mm^2^ dish, after one day cells were transfected with 1:1:1:1 ratio of the 3^rd^ generation lentiviral packaging plasmids (REV, RRE, VSV-G) and the Fast-FUCCI-puro plasmid (Addgene #86849), pCSII-CMV-MSC-IRES2-Bsr plasmid (CONT-EV) or RAB5Aoe plasmid pCSII-CMV-MSC-IRES2-Bsr-RAB5A (RAB5Aoe) which was generated via gibson assembly using primers fwd:*gcgctaccggtctcgagaattcATGGCTAGTCGAGGCGCAAC,* rev*:agaggggcggatccgcggccgcTTAGTTACTACAACACTGATTCCTGGTTG.* After two days, media was replaced with fresh DMEM supplemented with 10 mM sodium butyrate (Sigma, Cat. #B5887-1G) to induce virus production. On day 5, the supernatant was passed through a 0.45 µm filter (Millipore, Cat. #SLHV033RS) and supplemented with 8 µg ml^-1^ polybrene. One day prior to infection, 1 x 10^5^ KP or PC9 cells were seeded in a 6-well plate, the next day media was replaced with the filtered lentiviral suspension. Cells were expanded under 2 µg ml^-1^ Puromycin or 5 µg ml^-1^ Blasticidin selection and positive cells were sorted via FACS.

For the KP-pOncobow cells, a sorting strategy to was implemented to positively select for unrecombined cells expressing H2B-mTurquoise2 alone using single fluorescent protein expressing cells as controls (pPB-H2B-mTurquoise, pPB-yPET, pPB-dTomato, pPB-mKeima). Cells were trypsinized with 0.05% Trypsin-EDTA (Gibco, Cat. #25300120), washed twice with PBS, strained through a 40 µm cell strainer, and resuspended in PBS with 5 mM EDTA; 2% FBS at a concentration of 1 x 10^6^ cells ml^-1^. Cells were sorted on a BD FACS Aria III instrument with 445nm laser excitation source (BD Biosciences) for mTurquoise2. Following isolation of clonal populations, an imaging based 4-OHT recombination screen was conducted to ensure that selected clones recombined in the presence of low concentration 4-OHT (1-100 nM).

Cell lines (KP, PC9, A549, H1650, H1975, HCC827, CRUK0764, CRUK0879) were maintained in sub-confluent culture in the appropriate conditions. KP cells were cultured in DMEM; 20% FBS; 1% P/S, CRUK0764 and CRUK0879 cells were maintained in DMEM; 10% FBS;1% Insulin-transferrin-selenium (ITS) (Thermo Fisher Scientific, Cat. #41400-045); 1% P/S at 37 °C, 10% CO_2_. A549 cells were cultured in DMEM; 10% FBS; 1% P/S, PC9, HCC827, H1975 and H1650 cells were cultured in RPMI; 10% FBS; 1% P/S all at 37 °C, 5% CO_2_.

### Growth factors and inhibitors

pOncobow recombination was induced with 2-10 nM 4-hydroxy-tamoxifen (4-OHT) in DMSO. 10 µg human recombinant TGFβ was dissolved in ddH20 and applied in experiments at 1 ng ml^-1^. Blasticidin-S (Thermo Fisher, Cat. #R210-01) was diluted to a stock concentration of 10 mg m^l-1^ and applied at 5 µg ml^-1^. DMSO was used as a control at the appropriate concentration for all experiments where drug was applied.

### In vivo experiments

All mouse experiments were performed in compliance with the Francis Crick Institute Animal Welfare and Ethical Review Body and regulation by the UK Home Office project license PPL70/8380 and PP0736231. NSGTM mice (NOD.Cg-PrkdcscidIl2rgtm1Wjl/SzJ) used for the orthotopic lung tumour experiments were obtained from the Francis Crick core colony. The mice were housed in well ventilated cages at a constant temperature and humidity (23 °C ± 2 °C, 50– 60%) in a pathogen-free controlled environment, with a standard 12–12 h light–dark cycle, and food and water ad libitum.

### Orthotopic lung tumour models

KP-pOncobow or KP-FUCCI cells were trypsinized, washed in PBS, filtered through a 40 µm cell strainer and resuspended in PBS at a concentration of 1 x 10^4^ or 2 x 10^5^ cells ml^-1^ respectively. Cells were injected intravenously in 100 µl of PBS into the tail-vein of 8 to 12-week-old female NSG mice. The low concentration of cells injected was optimised to favour the spatial separation of tumours. Mice that received KP-pOncobow cells were given intraperitoneal injections of 4-hydroxy-tamoxifen (4-OHT) (Sigma Cat. #H7904-25MG) or ethanol control on days 3, 6, 10, or 15 following tail-vein injection. 4-OHT was dissolved in pure ethanol at a concentration of 20 mg ml^-1^ then diluted 1:1 in Kolliphor (Sigma, Cat. #C5135) to facilitate dissolution, reduce the viscosity and increase the ease of injection of 4-OHT (Chevalier, Nicolas and Petit, 2013). To achieve low density labelling, 4-OHT was diluted in sterile water at a final concentration of 4 mg ml^-1^ and mice received a dose of 20 µg g^-1^ body weight. Mice were monitored for changes in weight and breathing difficulties for 5-weeks or until the humane endpoint was reached. At endpoint, mice were euthanised via intraperitoneal overdose of pentobarbital or for perfusion via terminal anaesthetic.

Following confirmation of death, incisions were made to expose the trachea, diaphragm, heart and deflate the lungs. For *ex vivo* live cell imaging, lungs were immediately inflated by insertion of an 18 G cannula (BD Bioscience, Cat. #381347) into the trachea and injection of 1 ml of 2% Low melting point agarose (LMPA) (Sigma, Cat. #16520-050) dissolved in HBSS. For lung fixation at harvest, residual blood was removed via perfusion of the left ventricle of the heart with 10 ml of PBS followed by 5 ml Antigenfix (Solmedia, Cat. #P0016) to fix the lungs prior to inflation with 1 ml 1% LMPA dissolved in PBS. Following inflation, ice was placed around the thorax to solidify the agarose. The lungs were then resected, dissected into individual lobes, and processed for vibratome sectioning or optical projection tomography (OPT). Fixed lung lobes were incubated for 24 h in a solution of Draq5 (Sigma, Cat. #62251) diluted 1:100 in Antigenfix.

### Vibratome sectioning

To prepare for vibratome sectioning, individual lobes were washed in ice-cold HBSS then mounted in 2 or 8% LMPA to dampen the recoil of lungs when subjected to shear stress from the vibratome blade. A Leica Vibratome (Leica, Cat. #VT1200) affixed with a mounting blade (Ted Pella, Cat. # 121-4) was set to 0.8-1.5 mm s^-1^, 1-2 mm amplitude, section width of 200-300 µm, and serial sectioning mode. Tissue sections were collected in ice-cold HBSS once they floated from the main specimen mass. Live lung slices were then cultured in RPMI; 2% FCS; 1% P/S at 37°C, 5% CO_2_ prior to imaging. Fixed lung slices were stored in PBS at 4 °C until imaging.

### Ex vivo imaging

Live lung slices were immunostained with Alexa Fluor® 647 anti-mouse CD326 (Ep-CAM) Antibody (1:400) or Alexa Fluor® 647 anti-mouse/human CD324 (E-Cadherin) Antibody (1:400) (Biolegend) in culture media protected from light for 1h at 37 °C, 5% CO_2_. Then lung slices were mounted in 2% LMPA in a 12-well glass bottomed dish (MatTek) to ensure the tissue was flat against the coverslip and immobilised during imaging. Both live and fixed sections were imaged on an Olympus FV3000 confocal microscope (Olympus). For cell dynamics, images were taken every hour for 40 h within 24 h of lung resection. After imaging, lung slices were fixed in 4% PFA for 1 h at 20°C and stored in PBS at 4°C. For fixed lung slice imaging, slices were permeabilised for 30 min in PBS; 0.5% Triton X-100, stained with either Phalloidin–Atto 633 (0.02 nM) in PBS;0.1% Tween-20 for 1 h at 20 °C or with Click-IT Alexa Fluor647 EdU Staining Kit solution according to manufacturer’s instructions (Thermo-Fisher, Cat. #C10640) and then counterstained with 1 µg ml^-1^ DAPI in PBS (Sigma, Cat. #D9542) for 30 min at 20 °C. Cell tracking was performed using Imaris (9.9.1).

### Optical projection tomography

Optical projection tomography (OPT) was used for 3D imaging of KP tumours *ex vivo* in the context of the whole lung lobe environment. Here, individual lung lobes were mounted on a freely moving 360° rotating platform while a fixed-point laser captured the light signal and attenuation onto a camera sensor. First, resected lobes were embedded in 4% LMPA in a specially engineered mounting chamber. To minimize optical aberrations during imaging, the mounting chamber was polished with high grit sanding paper on the surfaces contacting LMPA. Briefly, lung lobes were skewered along the long axis with an 18 G needle, placed inside a single cylindrical mounting chamber, the end of the needle was affixed onto a Styrofoam dissection board under the mounting block and LMPA was added to encapsulate the lobe. Following solidification, the needle was removed and the lobe was taken out of the block.

The mounted lobes were then cleared using C_e_3D (Li, Germain and Gerner, 2017). C_e_3D was prepared by first diluting 37 °C N-methylacetamide (Sigma, Cat. #M26305) to 40% (v\ v) in PBS, then dissolving Histodenz (Sigma, Cat. #D2158) to 86% (w\v) in this diluent [1.45 g Histodenz per 1 ml 40 % N-methylacetamide] at 37°C to expedite dissolution. The final C_e_3D medium has been previously described to have a refractive index between 1.49 and 1.5 in range. Lungs mounted in agar were then submerged in 25 ml C_e_3D and cleared for at least 48 h but no more than 72 h. Immediately prior to imaging, a magnetized metal disk was superglued on top of the cleared agar block to facilitate attachment to OPT instrumentation.

The optical projection tomography (OPT) was undertaken using a custom designed instrument. This utilised a Zyla 5.5 acquisition camera (Andor Technology), a custom multimode laser-diode based excitation laser source (Tri-line, Cairn Research Ltd), a stepper motor (T-NM17A200, Zaber) and controller (X-MCB1, Zaber) to rotate the sample during image data acquisition, and a unity magnification telecentric lens (1.0 x SilverTL, Edmund Optics, Cat. #58-430). The sample was mounted in a glass cuvette filled with index-matching fluid and suspended via magnetic attachment to the stepper motor spindle, which was positioned laterally using linear stages (DTS25/M, Thorlabs), and the rotation axis tilt was adjusted using a kinematic mount (KM200, Thorlabs). Together, these degrees of adjustment enabled XYZ translation and tip/tilt of the sample in order to centre its position on the camera sensor, to centre it at the telecentric lens focal plane (or at an offset equivalent to half the sample thickness), and to arrange the rotation axis to be perpendicular to the optical axis of the imaging system and parallel to the columns of pixels on the sensor. This OPT set-up allowed provided tomographic 3D imaging with isotropic voxels of 13 µm^3^ on each side. OPT data acquisition was controlled using a custom OPT plug-in for *µManager* version 2.0γ (Edelstein *et al*., 2010). Optical filters were selected to match the fluorescent protein emission spectra of cells containing pOncobow and dyes from immunostaining samples prior to acquisition (See Table).

### Optical filters for OPT acquisition

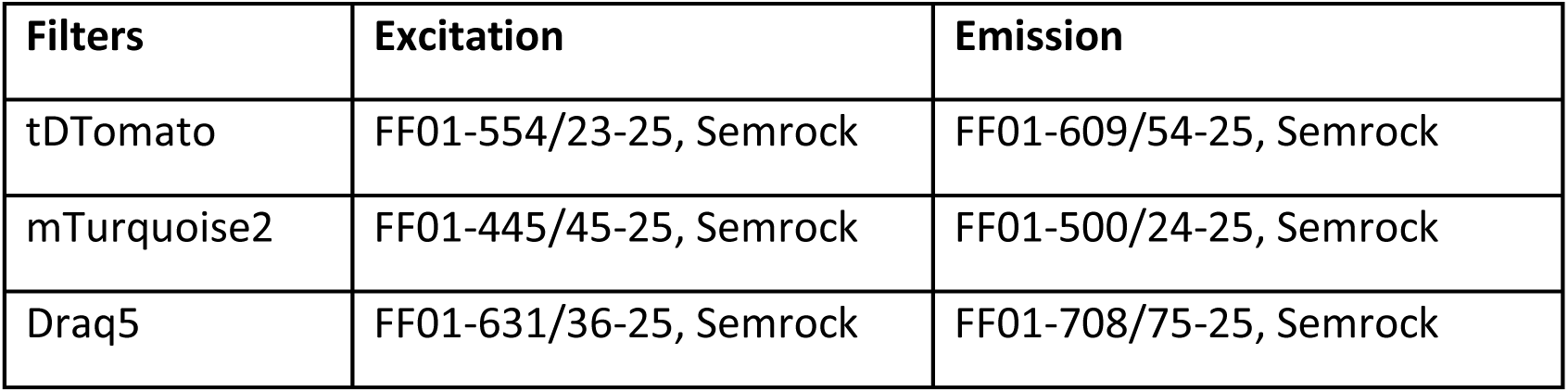

Samples were mounted in the glass cuvette and submerged with 25 ml Ce3D (previously used to clear the lung tissue samples). Projection images were acquired at 800 angles to allow adequate 3D reconstruction of the lung tissue samples bearing subclones. Following OPT image data acquisition, samples were equilibrated in PBS to allow for further analysis.

### 3D OPT image reconstruction and analysis

The OPT datasets comprising angular projections of the sample were reshaped as sinograms and 3D images were reconstructed with a Filtered Back Projection (FBP) algorithm using the standard *iradon* library function by MATLAB (The MathWorks Inc). The corresponding software and GUI is a part of ALYtools set of scripts (https://github.com/yalexand/ALYtools#ic_opttools). This MATLAB-based OPT image data reconstruction software is cross-OS capable and offers image operations acceleration via graphic processing unit (GPU) or multicore processing, as well as artefact correction and service options, such as axial mismatch correction, image downscaling, FBP setups control, I/O formats, batch processing, visualization of reconstructed volume, and more.

Subclone segmentation was performed using Icy with a combination of threshold-based segmentation, and size filtering to remove fluorescent artefacts present in each respective channel (De Chaumont *et al*., 2012). Following this process, segmented subclones (dTomato), clones (mTurquoise2), and lung (Draq5) were imported into a custom Python environment to render using Mayavi, an open-source toolkit for 3D rendering (Ramachandran and Varoquaux, 2010).

### Single cell-derived assays

Single cell-derived growth was ensured via plating using a CellenONE cell sorter. Following trypsinisation, cells were washed twice in PBS, strained through a 40 µm (KP) or 100 µm (human) cell strainer then resuspended in 2% FBS; 0.5 mM EDTA; PBS at concentration of 2 x 10^5^ cells ml^-1^.

### Colony assays and end-point imaging

Cells were single-cell plated in 96-well or at 10 cells well-^1^, in 24-well imaging plates (Ibidi Cat. #82426). After 3 days of growth, recombination of KP-pOncobow cells was induced with 2-10 nM 4-OHT for 6h. Growth factors or inhibitors were added on day 5. For therapy resistance experiments, 5 µg ml^-1^ blasticidin-S (Thermo Fisher, Cat. #R210-01) was added to the existing media on day 6. Due to differing growth rates and to ensure comparable colony sizes, all KP colonies were fixed on day 14 whereas human lung cell colonies were fixed on day 21 of growth. Prior to fixation, cells were incubated for 1 h with 10 µM EdU, then washed in PBS, fixed in 4% paraformaldehyde for 10 min at 20 °C, washed three times in PBS and stored at 4 °C. Colonies were stained with Click-iT AlexaFluor647 EdU Staining Kit according to manufacturer’s instructions and nuclei were counterstained with 1 µg ml^-1^ DAPI. Fixed colonies were imaged using an Olympus FV3000 confocal microscope.

### Colony ‘re-growth’ assays

For the cloning ring experiments, single KP-pOncobow cells were plated using the CellenONE into 24-well plates containing a 15.6 mm growth surface (Ibidi, Cat.#82406). Media was replaced after 15 days of growth, then on day 25 the media was removed and a 3.2 mm diameter cloning disk (Sigma, Cat. #Z374431) was pushed down on the centre of the colony for 1 min to remove the cells underneath. Following this, media was replaced, and cells were grown for another 3 days. Colony regrowth was then imaged using a confocal microscope.

For hole experiments, KP-FUCCI or KP-pOncobow cells were plated at a density of 5 cell well^-1^ in a 24-well imaging plate (Ibidi, Cat #82425). Recombination was induced in KP-pOncobow colonies on day 7 of growth via treatment with 2 nM 4-OHT for 6 h. On day 12 for KP-FUCCI or day 15 for KP-pOncobow, colonies were imaged using a Nikon Ti2 inverted microscope prior to hole generation. Then the centre of the colony was removed using a P200 pipette tip to scratch away cells. Colonies were then washed with PBS and new media was added. Then Colony regrowth was imaged every 120 min for 96 h for KP-FUCCI colonies or every 120-240 min for 96 h for KP-pOncobow colonies.

### Collagen ‘crash’ assay

Collagen gels were prepared from type I collagen (Corning #354249). Prior to experiments, the collagen I solution was neutralized with 0.1 M NaOH to pH = 7, diluted to a final concentration of 8 mg ml^-1^, mixed with 2 μm fluorescent beads as fiduciary markers, and then poured into culture-insert 2 well chambers (Ibidi GmbH; Germany, Cat. #80209-150) placed onto glass bottom dishes. Following collagen polymerisation at 37 °C, KP cells expressing GFP were then seeded around the inserts. Upon reaching confluency, the inserts were removed, causing KP cells to migrate toward collagen gels. The collision of KP cells with the collagen and its subsequent deformation was imaged using confocal microscopy.

### Live cell imaging

All live cell imaging was performed on a Nikon Ti2 inverted microscope with perfect focus system, teledyne-photometrics Prime BSI scientific CMOS camera, an ASI motorised XY stage with piezo Z, and an Okolab environmental chamber and CO_2_ mixer. For dynamic imaging, colonies were imaged on day 5, 12 and 19 of growth every 6-10 min using a Plan Apochromat 20x 0.75NA Ph2 objective. For imaging of colony expansion, colonies were imaged using a Plan Fluor 10x 0.3NA Ph1 objective every 2 h from day 5 (KP-FUCCI, KP-pOncobow) or day 6 (KP-pOncobow-BSR) until day 12 of growth. Acquisition software was custom designed based on a build of µManager software (Edelstein *et al*., 2010). Tiled colony images with 10% overlap were reconstructed using custom FIJI scripts (Preibisch, Saalfeld and Tomancak, 2009; Schindelin *et al*., 2012).

### Single colour colony growth rate

To establish growth rate of expanding subclones, KP cells expressing single colour constructs (pPB-H2B-mTurquoise, pPB-yPET, pPB-dTomato) were plated at density of 1.5 x 10^3^ cell well^-1^ in a 96-well plate. Cells were left to adhere overnight then placed in an Incucyte Zoom (Essen Biosciences) and imaged every 3 h for 7 days. To analyse growth rate, confluency estimates were generated from the phase images using the Incucyte software according to manufacturer instructions. Logistic growth curves were fitted to allow comparison of growth rates between different the different coloured populations.

### CAF co-culture experiments

Human CAF primary cell cultures were established through the Tracking Cancer Evolution through Therapy (TRACERx) clinical cohort study (REC reference 13/LO/1546). CAF cultures were initiated as explant cultures of tumour tissue from patients with lung adenocarcinoma in DMEM containing 10% FBS and 1% P/S. The fibroblast identity of the resulting cells was confirmed by immunostaining before cultures were immortalised using hTERT (described in Gaggioli et al 2007). KP-pOncobow cells were single-cell plated at 10 cells well-^1^, in 24-well imaging plates and were grown for 3 days. Recombination of KP-pOncobow cells was induced with 2 nM 4-OHT for 6 h, then colonies were washed with PBS and new media was added. CAFs were added after 4-OHT treatment, on day 3 or 5 of colony growth at a density of 4x10^4^ -1.6x10^5^ cells well^-1^ to form a confluent layer around the KP-pOncobow colonies. Media was changed on day 10 of the assay.

### Immunofluorescence and immunoblotting

Following fixation, cells were blocked for 1 h at 20 °C in blocking buffer (2% BSA-5% FBS;0.1% Tween-20-PBS). Primary antibodies (1:250-1:400) in staining buffer (1% BSA;2.5% FBS;0.1% Tween-20-PBS) were incubated overnight at 4 °C. Following three 5 min washes in 0.1% Tween-PBS, secondary antibodies (1:1000) were incubated for 1 h at 20 °C, then washed twice with 0.1% Tween-PBS, counterstained with 1 µg ml^-1^ DAPI, washed twice in PBS and imaged. For immunoblotting, cells were lysed in ice-cold RIPA buffer (20 mM Tris-HCl pH7.4;150 mM NaCl; 1% NP40; 0.1% Sodium deoxycholate; 0.1% SDS) with PhosStop and Complete inhibitors, protein concentration determined using BCA and 30 µg protein separated by SDS-PAGE, transferred to a PVDF membrane, blocked with 5% BSA; TBS; 0.1% Tween-20 at 4 °C overnight. Membranes were probed with primary and secondary antibodies (1:1000-1:10,000) for 1 h washed in TBS; 0.1% Tween-20 and detected using chemiluminescence. See table for individual antibody details and dilutions.

### Quantitative analysis

Quantitative analysis included image segmentation and tracking using FIJI (v1.53t), quantification using custom R scripts and Python Shapely, statistical analysis and plotting using R. For colony end-point experiments segmentation was automated using custom FIJI scripts (Schindelin *et al*., 2012). Briefly, colony area was segmented using the DAPI channel via size 50 scaled radius gaussian blur and manual thresholding, then subclone segmentation was performed using a size 3 scaled radius gaussian blur and threshold values optimised for each colour fluorescent protein subtracting any dual-positive subclones based on area overlap. Threshold values for colony area and subclone area were optimised for each experiment to account for variation during image acquisition or protein expression levels. Area was quantified from the coordinates using the Shapely package (1.8.0) in python (Gillies, 2007) where subclones were defined as contiguous colour patches with an area greater than 100 µm^2^. Colonies with less than one subclone or a subclone occupying greater than 30% the colony area were excluded to account for low recombination efficiency or limit the likelihood of calling colour patches that contained adjacent subclones expressing the same colour. EdU and DAPI intensity values were used for analysis of proliferation in colonies and for density experiments EdU and DAPI maxima counts were extracted using optimised thresholds. Nuclear area was quantified via segmentation with StarDist using default model parameters (Schmidt *et al*., 2018) filtering nuclei below 170 µm^2^ (max non-mitotic area) to avoid quantifying cells in mitosis when the hGEM-AG signal becomes cytoplasmic. For cellular dynamics, cells were tracked based on nuclear segmentation from timelapse images taken every 6 or 10 min for 18 h using StarDist-TrackMate with default model parameters and track splitting and merging disabled (Ershov *et al*., 2022). Cell mixing analysis involved using points from individual tracks to construct a convex hull with SciPy (1.10.1) then counting the number of tracks intersecting with the hull-defined area.

### Reagent Availability

The biological reagents described above are either available from the sources indicated or upon request from

### Statistics

Statistical analysis was performed in R (4.0.3 or 4.2.2) using base R Welch unpaired two-sample t.test() function for comparison between conditions in analysis of readouts from simulated colonies and patient data or rstatix (v0.7.0) functions wilcox_test() for two-sample pairwise comparisons or kruskal_test() and post-hoc dunn_test() for multiple comparisons with Benjamin-Hochberg p-value correction for experimental data. All tests were two-tailed. When applicable, the n, p-value, t-value, and degrees of freedom are indicated in figure legends, p-values are shown to one significant figure as ‘ns’ p > 0.05, * p < 0.05, ** p < 0.01, *** p < 0.001, **** p < 0.0001. All plots were generated with ggplot2 (3.3.6 or 3.4.1) Matpotlib (5.3.2) or Plotly (5.7.0). All boxplots show median line, interquartile range and error bars are mean ± standard deviation, if present mean is shown as a red dot. All violin plots show median line.

## Data Availability

Representative movies of experiments and simulations are available as Supplementary Videos alongside the main manuscript. The data frames of source data for generating results reported in the manuscript will be available to editors and referees upon request and will be publicly available on figshare repository at publication.

Patient data (the RNA-seq and whole-exome sequencing data) from the TRACERx Lung study are published in Frankell AM, Dietzen M, Al Bakir M, Lim EL, Karasaki T, Ward S, Veeriah S, Colliver E, Huebner A, Bunkum A, Hill MS, Grigoriadis K, Moore DA, et al. The evolution of lung cancer and impact of subclonal selection in TRACERx. Nature. doi.org/10.1038/s41586-023-05783-5 In press April 12th 2023.

The RNA-seq and whole-exome sequencing data (in each case from the TRACERx study) used during this study have been deposited at the European Genome–phenome Archive, which is hosted by the European Bioinformatics Institute and the Centre for Genomic Regulation, under the accession codes EGAS00001006517 (RNA-seq) and EGAS00001006494 (whole-exome sequencing). Access is controlled by the TRACERx data access committee. Details on how to apply for access are available at the linked page.

## Code Availability

Custom computer code for the agent-based model and analysis scripts for generating results reported in the manuscript will be available to editors and referees upon request and will be publicly available on GitHub repository at publication.

## Movie Legends

**Movie 1:** Optical projection tomography 3D reconstruction of whole lung lobe bearing KP-Oncobow tumours. Recombination was induced with 4-OHT on day 3 post tail-vein injection into NSG mice, mTurquoise and dTomato subclones are shown in blue and red respectively with Draq5 counterstain in grey. Scale bar = 1000 µm.

**Movie 2:** Representative time lapse movies of KP-pOncobow and KP-FUCCI single cell colony expansions every 2 h for 72 h starting on day 7 (168 h) of colony growth. For the KP-pOncobow colony, colours represent subclones H2B-mTurquoise (cyan), yPET (yellow), dTomato (red). mKeima subclones could not be imaged due to the dichroic set-up of the microscope but the uncoloured subclone patches are predicted to express mKeima. For the KP-FUCCI colony, Geminin-Azami Green (mAG-hGEM) is shown in green and Cdt1-Kuabira-Orange (mKO-CDT1) is shown in magenta. Scale bar = 500 μm.

**Movie 3:** Representative time lapse movies of central and edge regions from a KP-FUCCI single cell colony imaged every 6 min for 18 h on day 12 of colony growth. Geminin-Azami Green (mAG-hGEM) is shown in green and cdt1-Kuabira-Orange (mKO-CDT1) is shown in magenta. Scale bar = 100µm.

**Movie 4**: Time evolution of proliferation and subclones in representative simulations with varying model settings of cell random motility and drag coefficient. As indicated in the movie, simulations with “Reference”, “Intrinsic motility”, and “Low drag coef.” model settings are sequentially shown.

**Movie 5:** Representative time lapse movies of central regions from control and TGFβ treated KP-FUCCI single cell colonies imaged every 6 min for 18 h on day 12 of colony growth. The summed intensity for the Geminin-Azami Green and cdt1-Kuabira-Orange is shown in grey. Scale bar = 100µm.

**Movie 6**: Time evolution of therapy-resistant cells in representative simulations with varying model settings of cell random motility. As indicated in the movie, simulations with model settings low and high random cell motility, namely, “Reference” and “Intrinsic motility”, are sequentially shown.

**Movie 7:** Representative time lapse images of control, control + 5 µg ml^-1^ Blasticidin, TGFβ and TGFβ + 5 µg ml^-1^ Blasticidin (blasti) KP-pOncobow-BSR colonies every 2 h from day 6 (144 h) to day 10 of growth. Colours represent subclones H2B-mTurquoise (cyan), yPET-BSR (yellow), dTomato (red). Scale bar = 500 μm.

## Acknowledgements

We are grateful to David Novo and the Francis Crick Institute Science Technology Platforms, especially Advanced Light Microscopy, Flow Cytometry, Biological Research Facility, Cell Services and Scientific Computing, for their scientific and technical support.

We thank colleagues in Tumour Cell Biology Laboratory for their constructive feedback on this project over the years. We thank Karin Schlegelmilch and Robert Jenkins for the critical and helpful comments on the manuscript. We thank Nicholas Mcgranahan, Rachel Rosenthal, Carlos Martinez Ruiz, Kristiana Grigoriadis, Oriol Pich, David Moore, and Takahiro Karasaki for the insightful discussions on patient data in the TRACERx Lung study.

This work was funded by the Francis Crick Institute which receives its core funding from Cancer Research UK (FC001144, FC001003), the UK Medical Research Council (FC001144, FC001003), and the Wellcome Trust (FC001144, FC001003).

X.F. and E.S. were additionally supported by ERC Advanced Grant CAN_ORGANISE, Grant agreement number 101019366. Y.N was supported by a Grant-in-aid for JSPS Overseas Research Fellowship (No. 201860634). R.E.H. was supported by a Sir Henry Wellcome fellowship (WT209199/Z/17/Z). D.B. was supported by funding from a Cancer Research UK (CRUK) Early Detection and Diagnosis Project award, the Idea to Innovation (i2i) Crick translation scheme supported by the Medical Research Council, the National Institute for Health Research Biomedical Research Centre and the Breast Cancer Research Foundation (BCRF). A.L.M receives post-doctoral funding from AstraZeneca. Y.A. receives funding from CRUK Accelerator grant (A29368). C.S. is a Royal Society Napier Research Professor (RSRP\R\210001). His work is supported by the Francis Crick Institute that receives its core funding from Cancer Research UK (CC2041), the UK Medical Research Council (CC2041), and the Wellcome Trust (CC2041). For the purpose of Open Access, the author has applied a CC BY public copyright licence to any Author Accepted Manuscript version arising from this submission. C.S. is funded by Cancer Research UK (TRACERx (C11496/A17786), PEACE (C416/A21999) and CRUK Cancer Immunotherapy Catalyst Network); Cancer Research UK Lung Cancer Centre of Excellence (C11496/A30025); the Rosetrees Trust, Butterfield and Stoneygate Trusts; NovoNordisk Foundation (ID16584); Royal Society Professorship Enhancement Award (RP/EA/180007); National Institute for Health Research (NIHR) University College London Hospitals Biomedical Research Centre; the Cancer Research UK-University College London Centre; Experimental Cancer Medicine Centre; the Breast Cancer Research Foundation (US) (BCRF-22-157); Cancer Research UK Early Detection an Diagnosis Primer Award (Grant EDDPMA-Nov21/100034); and The Mark Foundation for Cancer Research Aspire Award (Grant 21-029-ASP). This work was supported by a Stand Up To Cancer-LUNGevity-American Lung Association Lung Cancer Interception Dream Team Translational Research Grant (Grant Number: SU2C-AACR-DT23-17 to S.M. Dubinett and A.E. Spira). Stand Up To Cancer is a division of the Entertainment Industry Foundation. Research grants are administered by the American Association for Cancer Research, the Scientific Partner of SU2C. CS is in receipt of an ERC Advanced Grant (PROTEUS) from the European Research Council under the European Union’s Horizon 2020 research and innovation programme (grant agreement no. 835297). The TRACERx study is funded by Cancer Research UK (CRUK; C11496/A17786) and the derivation of TRACERx patient models was supported by the CRUK Lung Cancer Centre of Excellence and a Sir Henry Wellcome fellowship (to R.E.H.; WT209199/Z/17/Z).

## Author Contributions

Conceptualisation and study design: A.B., X.F., S.B., E.S.

Acquisition of data: A.B., X.F., S.B., Y.N., H.M., R.E.H., S.H., C.S.

Development of methodology: A.B., X.F., S.B., S.H., D.B., A.L.M., S.K., Y.A., P.F., J.M., P.A.B., C.S, E.S.

Analysis and interpretation of data: A.B., X.F., S.B., E.S.

Writing and editing: X.F., S.B., A.B., E.S. with inputs from all authors

## Materials & Correspondence

Correspondence and material requests should be addressed to Xiao Fu and Erik Sahai.

---

**Supp. Figure 1:**
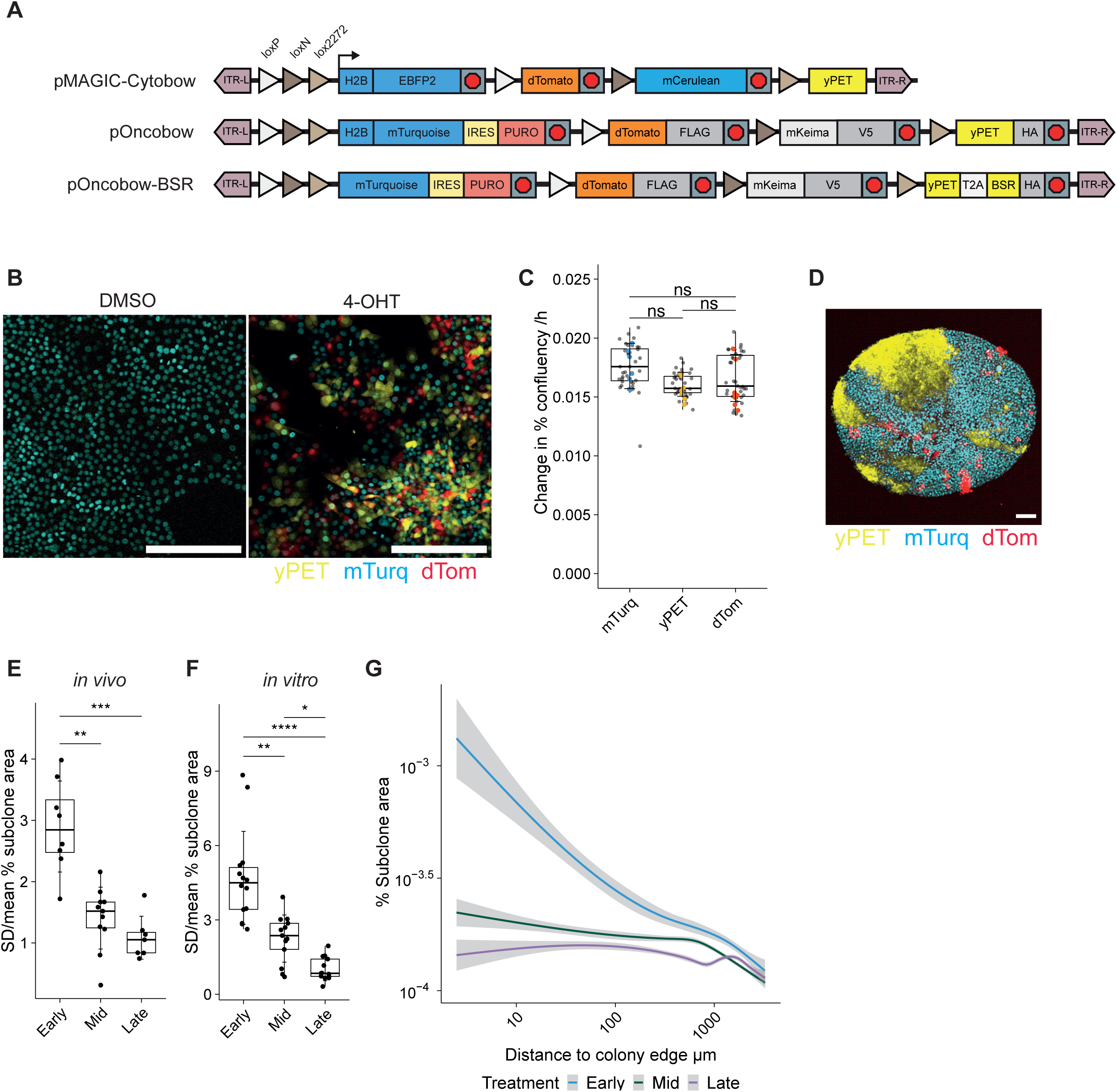
Multi-colour lineage tracing reveals unequal expansion of neutral subclones. A. Schematic of full cassette structure for pMAGIC-Cytobow (i), pOncobow (ii) and pOncobow-BSR (iii). B. Representative images of recombination in KPCreERT^2^ (KP) pOncobow cells after 6 h, 20 nM 4-OHT treatment or DMSO control. Scale bar = 500 µm. C. Quantification of the growth rate of KP cells expressing single colour constructs (pPB-H2B-mTurquoise, pPB-yPET, pPB-dTomato). Logistic growth curves were fitted to confluency estimates based on phase contrast images taken on an Incucyte Zoom taken every 3 h for 7 days to allow comparison of growth rates between different the different coloured populations. N = 7 replicates. D. KP-pOncobow spheroid displaying mTurquoise, dTomato and yPET subclones shown in the xy plane. Scale bar = 100 µm. Spheroids were generated by hanging drop method (Conti et al., 2020) and recombination was induced with 4-OHT. E. Quantification of the standard deviation over the mean of subclone area normalised to tumour area for each labelling time. N = 2 independent experiments, early n = 8, mid n = 11, late n = 7 tumours. F. Quantification of the standard deviation of over the mean subclone area normalised to colony area for each labelling time. N = 2 independent experiments, early n = 14, mid n = 13, late n = 13 colonies. G. Quantification of subclone area normalised to colony area versus subclone normalised minimum distance to colony edge for each labelling time point. N = 2 independent experiments, early n = 14, mid n = 13, late n = 13 colonies. Subclone area was defined as contiguous single colour patches greater than a minimum cell area of 100 µm^2^. All error bars are mean ± sd, pairwise comparisons by Dunn’s test with Benjamin-Hochberg p-value correction, ns p > 0.05. * p < 0.05, ** p < 0.01, *** p < 0.001, **** p < 0.0001.

**Supp. Figure 2:**
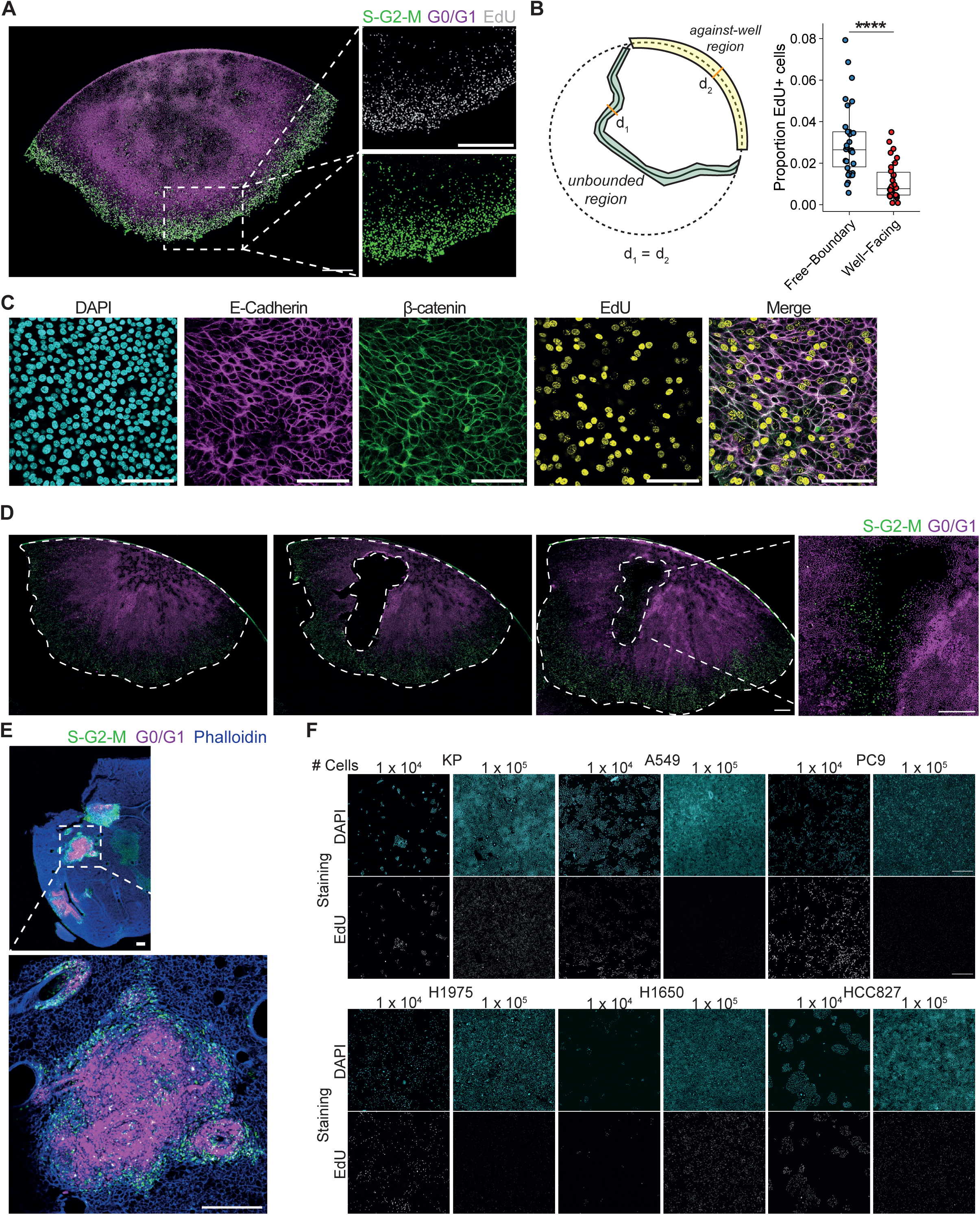
Analysis of spatial growth patterns. A. Representative image of KP-FUCCI single cell colony following proliferative arrest after 14 days of growth, ROIs highlight patterns of cell proliferation for S-G2-M (green) and S phase (EdU) labels. N = 3 independent experiments, scale bar = 500 μm. B. Schematic showing regions of colony periphery quantified and the proportion of EdU+ nuclei at the colony periphery facing the well or the free boundary. C. Representative immunofluorescence staining of DAPI, E-Cadherin and EdU incorporation in KP control colony at the boundary. N = 3 independent experiments, scale bar = 500 μm. D. Representative images of KP-FUCCI single cell colony grown for 12 days (pre-hole) then immediately after a patch of cells was removed at the colony centre (post-hole) and after 48 h of growth (48 h post hole). Inset is zoom of central region 48 h post hole creation, white dashed lines denote colony periphery and region removed. Scale bar = 500 µm, N = 3 experiments, n = 5 colonies. E. Representative image of whole lung slice and inset of KP-FUCCI tumour from an *ex vivo* lung slice. Lungs were harvested from NSG mice on day 25 post tail vein injection and sliced into 300 µm sections using a vibratome. Lung slices were fixed in 4% PFA, stained with phalloidin and DAPI then imaged. N = 2 independent experiments, n = 8 mice, scale bar = 500 μm. F. A549, PC9, H1975, H1650 and HCC827 human lung cancer cell lines were plated at 1 x 10^4^, 2.5 x 10^4^, 5 x 10^4^ and 1 x 10^5^ cells per well, incubated for 48-72 h. A 1 h 10 μM EdU pulse was performed prior to fixation and immunostaining for DAPI and EdU. Cells were imaged and the number of EdU+ and DAPI+ nuclei quantified. Representative images for DAPI and EdU staining of lung cancer cell lines at low and high density. Scale bar = 250 μm.

**Supp. Figure 3:**
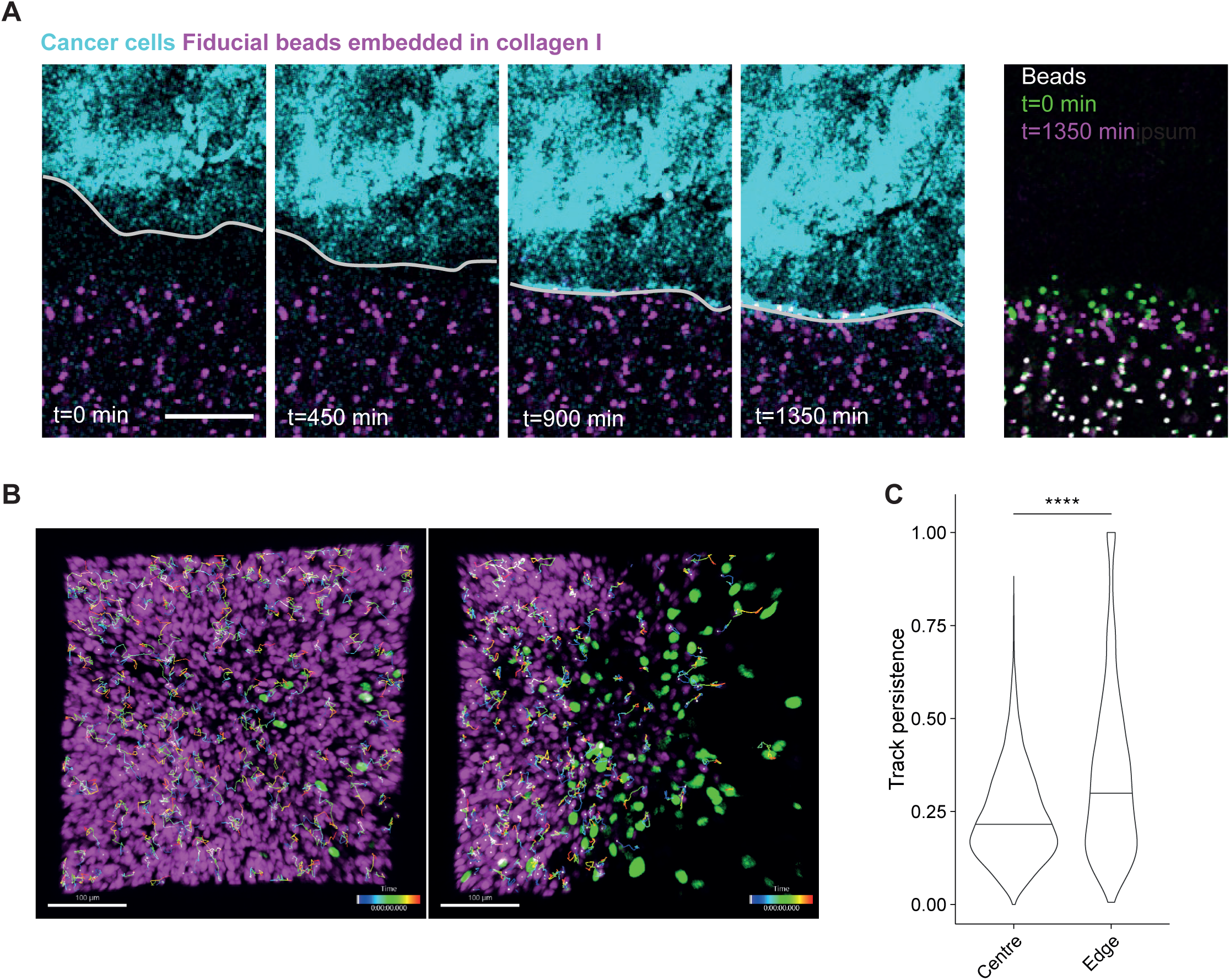
Analysis of migratory dynamics and forces. A. Images showing KP cells (cyan) moving towards and colliding with a 3 mg ml^-1^ collagen gel embedded with fluorescent beads (magenta). Right most panel shows the displacement of the beads over a 450 min period with starting position shown in green and final position in magenta. One representative experiment of 3 shown. Scale bar = 200 μm. B. Representative images of cell tracks in central and edge regions of a KP-FUCCI tumour from an *ex vivo* lung slice. Lungs were harvested from NSG mice on day 25 post tail vein injection and sliced into 300 µm sections using a vibratome. Lung slices were mounted in agarose and imaged every hour for 40 hrs for cell tracking. Scale bar = 100 µm C. Quantification of track persistence for cells in central and edge regions of *ex vivo* tumours performed using Imaris. Median shown as horizontal line, error bars are mean ± sd, pairwise comparisons by Wilcoxon signed rank test, *** p < 0.001, N = 4 mice, centre n = 4364, edge n = 2961 cells.

**Supp. Figure 4:**
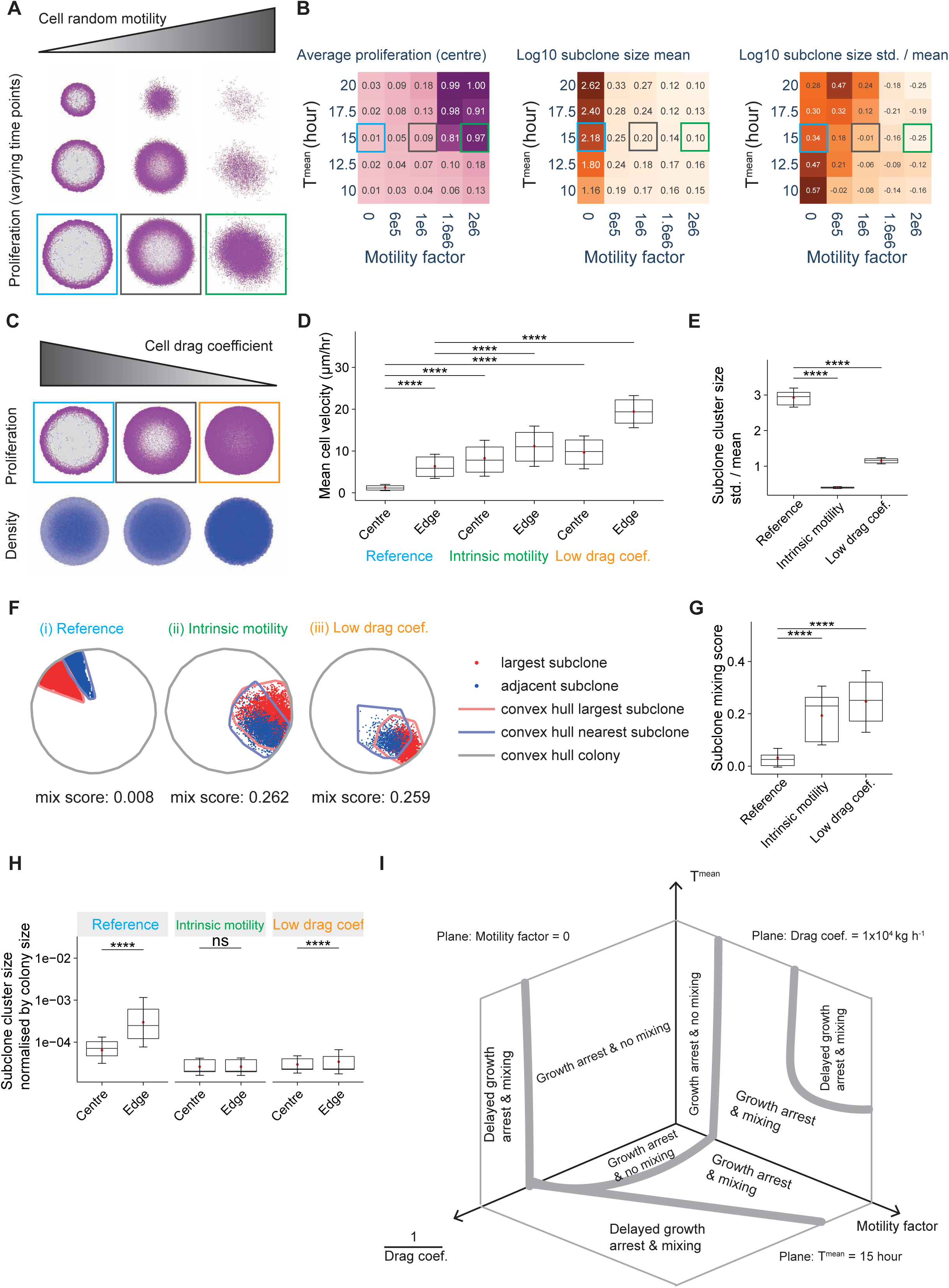
Computational modelling predicts cell migratory dynamics alters subclone evolutionary fates. A. Spatial patterns of proliferation in representative simulations with varying levels of random cell motility. Proliferatively active and arrested cells are labelled in magenta and light grey, respectively, with patterns at multiple time points (colony size of 2, 3, and 4 mm, respectively). Each column reflects snapshots from the same representative simulation. Boxes in different colours in (A) and (B) correspond to varying model settings of random cell motility. B. Phase diagrams of average fraction of proliferative cells at the colony centre, Log10-transformed average subclone cluster size, and Log10-transformed std./mean of subclone cluster sizes, with respect to intrinsic doubling time (𝑻^𝒎ean^) and random cell motility. N = 16 simulations for each parameter combination. C. Spatial patterns of proliferation and cell density in representative simulations with varying levels of cell drag coefficient. Proliferatively active and arrested cells are labelled in magenta and light grey, respectively, with patterns at multiple time points. Local cell density is labelled in blue, with greater intensity indicating higher density. Each column reflects snapshots from the same representative simulation. Boxes in different colours correspond to varying model settings of drag coefficient. Simulated colonies in these representative simulations have a size of about 4 mm. D. Mean cell velocity over a duration of 18 hours, at the colony centre versus edge, in simulations with different model settings of cell random motility and drag coefficient. N = 16 simulations for each condition. Welch two-sample t-test is performed. “****” reflects p <0.0001. For comparisons from left to right, t-values and degrees of freedom, (t, df), are: (-87.331, 2970), (-79.672, 2587.7), (-111.6, 3028.1), (-42.743, 3967.3), (-142.66, 5219). E. The standard deviation of subclone cluster size divided by the mean, compared between different model settings of cell random motility and drag coefficient. The subclone cluster size is measured as the number of cells in the subclone cluster divided by the total number of cells in the colony. In contrast to the Main Figure 4I, a subclone is defined as a spatially contiguous patch of cells with an identical lineage (see Supplementary Methods). N = 16 simulations for each condition. Welch two-sample t-test is performed. “****” reflects p <0.0001. For comparisons from left to right, t-values and degrees of freedom, (t, df), are: (37.693, 15.286), (25.265, 18.141). F. Convex hull demarcation of the largest subclone (red line) and one adjacent subclone (green line). Convex hull demarcation of the colony is also shown (grey line). Subclone mixing score is calculated as the fraction of all cells within red convex hull being in green. G. Subclone mixing scores in simulations with varying conditions of cell random motility and drag coefficient. N = 16 simulations for each parameter condition. Welch two-sample t-test is performed. “****” reflects p <0.0001. For comparisons from left to right, t-values and degrees of freedom, (t, df), are: (-5.4653, 17.999), (-6.9808, 17.736). H. Subclone cluster size normalised by colony size, at the colony centre versus edge, in simulations with different model settings of cell random motility and drag coefficient. N = 16 simulations for each condition. Welch two-sample t-test is performed. “****” reflects p<0.0001; “ns” reflects not significant. For comparisons from left to right, t-values and degrees of freedom, (t, df), are: (-15.614, 1527.1), (-0.79089, 52434), (-23.101, 28443). I. A schematic illustration summarising states of growth arrest and subclone mixing with respect to the intrinsic doubling time (𝑻^𝒎ean^), cell random motility, and drag coefficient. In (D), (E), (G), and (H), two ends of the box reflect the lower and upper quartiles, respectively, and the horizontal line dividing the box reflects the median. The red dot and error bar indicate the mean and the standard deviation, respectively.

**Supp. Figure 5:**
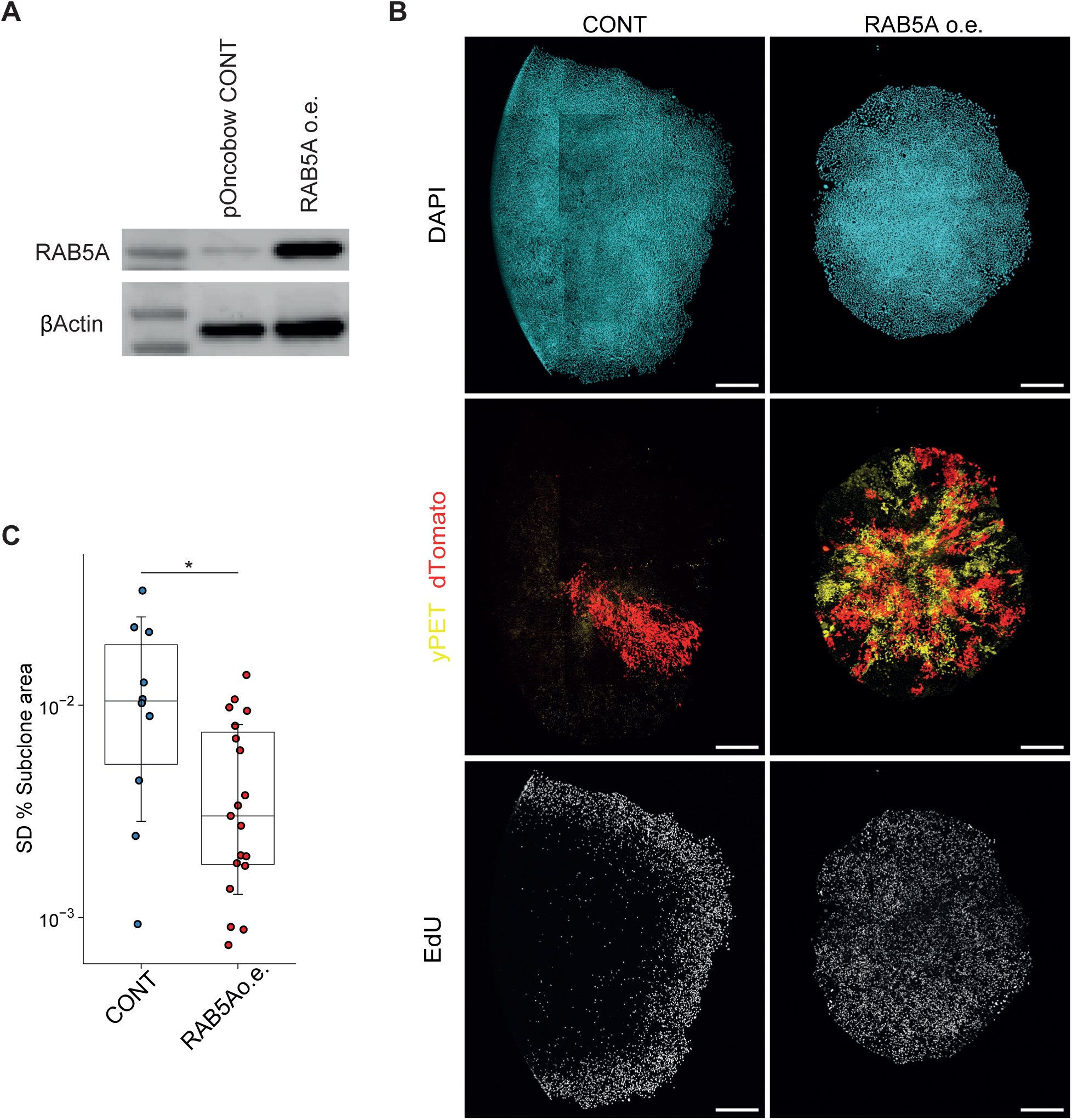
Cell migration alters subclonal fates. A. Western blot showing overexpression of RAB5A (RAB5Ao.e.). B. Representative confocal images of KP-pOncobow empty vector control (CONT) and RAB5A overexpressing (RAB5Ao.e.) colony subclone patterns colours represent DAPI (cyan), yPET (yellow) and dTomato (red) subclones, EdU (grey). Single KP-pOncobow cells overexpressing RAB5Aoe or CONT-EV were plated, treated with 2 nM 4-OHT for 6 h on day 3. Cells were then left in culture until day 14 when a 1 h 10 μM EdU pulse was performed prior to fixation and immunostaining for DAPI and EdU. Scale bar = 500 μm C. Quantification of standard deviation of normalised subclone size. Error bars are mean +/- standard deviation, pairwise comparisons by Wilcoxon signed rank test, * p < 0.05, n = 22 colonies, N = 2 independent experiments.

**Supp. Figure 6:**
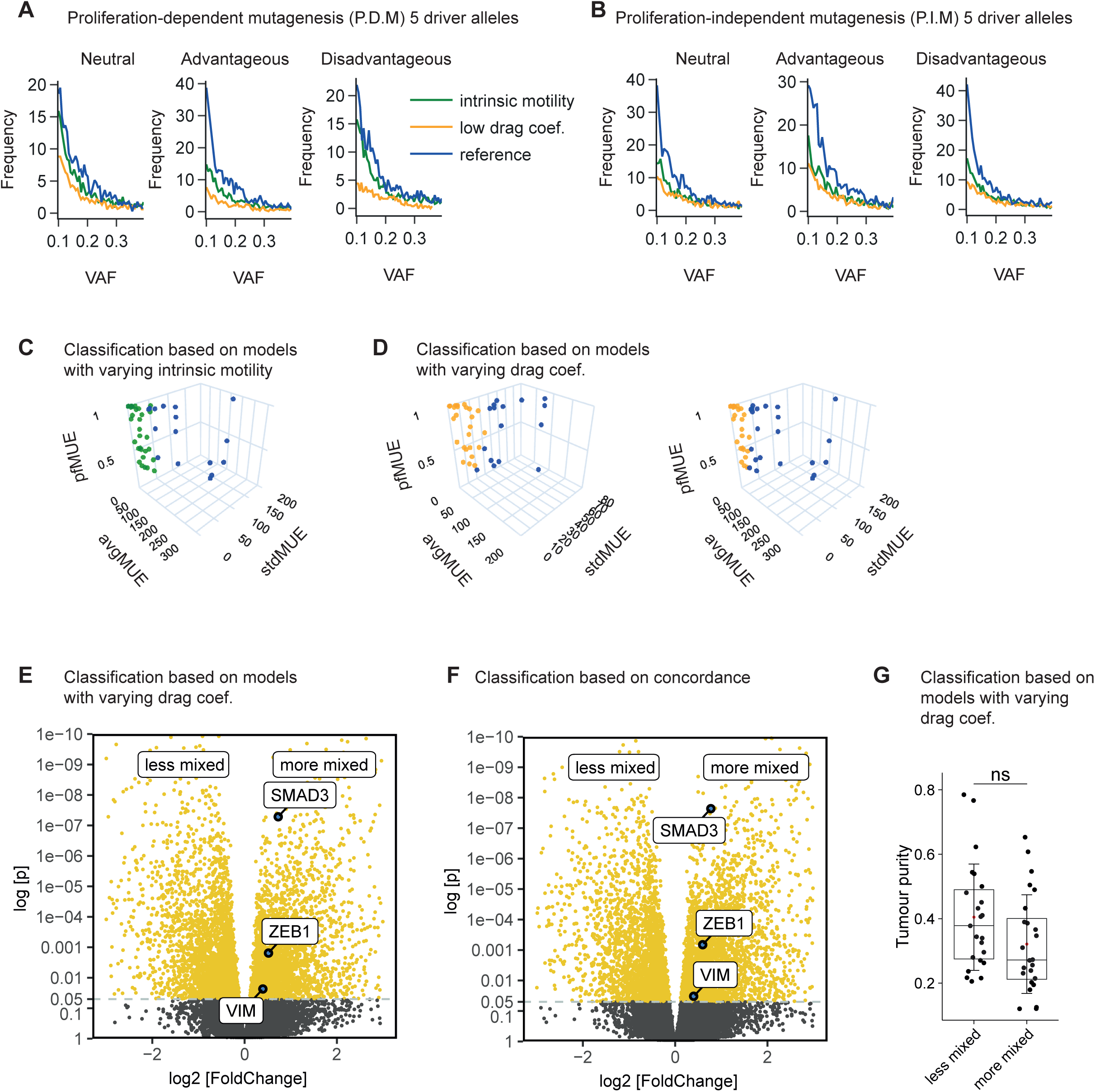
Computational modelling links TGFβ and EMT with subclonal mixing in human lung cancer. A. Frequency distributions of regional variant allele frequency (VAF) in simulations with varying model settings of random cell motility and drag coefficient, labelled as “Reference” (blue), “Intrinsic motility” (green), and “Low drag coef.” (orange) respectively, grouped into subsets of simulations with different evolutionary conditions. Lines reflect frequency distribution of VAFs in simulated tumours, averaged across regions of N = 40 simulations per condition. This set of simulations were based on implementation of proliferation-dependent mutational processes (“P.D.M”). B. Description as (A). This set of simulations were based on implementation of proliferation-independent mutational processes (“P.I.M”). C. Inferred mixedness of LUAD tumours with predominantly solid histological subtype in the TRACERx Lung study plotted in the VAF feature space, based on a Support Vector Machine (SVM) classifier trained using simulations with low vs high levels of random cell motility. Tumours with more extreme MUE features that are not shown in Main Figure 6F are presented in this plot. D. Inferred mixedness of LUAD tumours with predominantly solid histological subtype in the TRACERx Lung 421 study plotted in the VAF feature space, based on a SVM classifier trained using simulations with low vs high levels of drag coefficient. Tumours with more extreme MUE features are presented only in the right panel (Also refer to Supplementary Table 1). E. Differential expression of genes between subsets of tumours inferred to be “more mixed” (N = 61 regions from 20 tumours) and “less mixed” (N = 60 regions from 20 tumours). Select significantly differentially expressed EMT genes are highlighted in the plot. Inference of mixedness was based on a SVM classifier trained using simulations with low vs high levels of drag coefficient. F. Differential expression of genes between subsets of tumours inferred to be “more mixed” (N = 61 regions from 20 tumours) and “less mixed” (N = 47 regions from 15 tumours). Select significantly differentially expressed EMT genes are highlighted in the plot. Inference of mixedness was based on concordant classifications between motility- and drag-based SVM classifiers. G. Tumour purity in subsets of tumours inferred to be “less mixed” (N = 23 tumours) and “more mixed” (N = 24 tumours), respectively. Inference of mixedness was based on a SVM classifier trained using simulations with low vs high levels of drag coefficient. Welch two-sample t-test is performed. “ns” reflects not significant, p = 0.07899. The t-value and degree of freedom, (t, df), is: (1.7979, 44.388). In (G), two ends of the box reflect the lower and upper quartiles, respectively, and the horizontal line dividing the box reflects the median. The red dot and error bar indicate the mean and the standard deviation, respectively.

**Supp. Figure 7:**
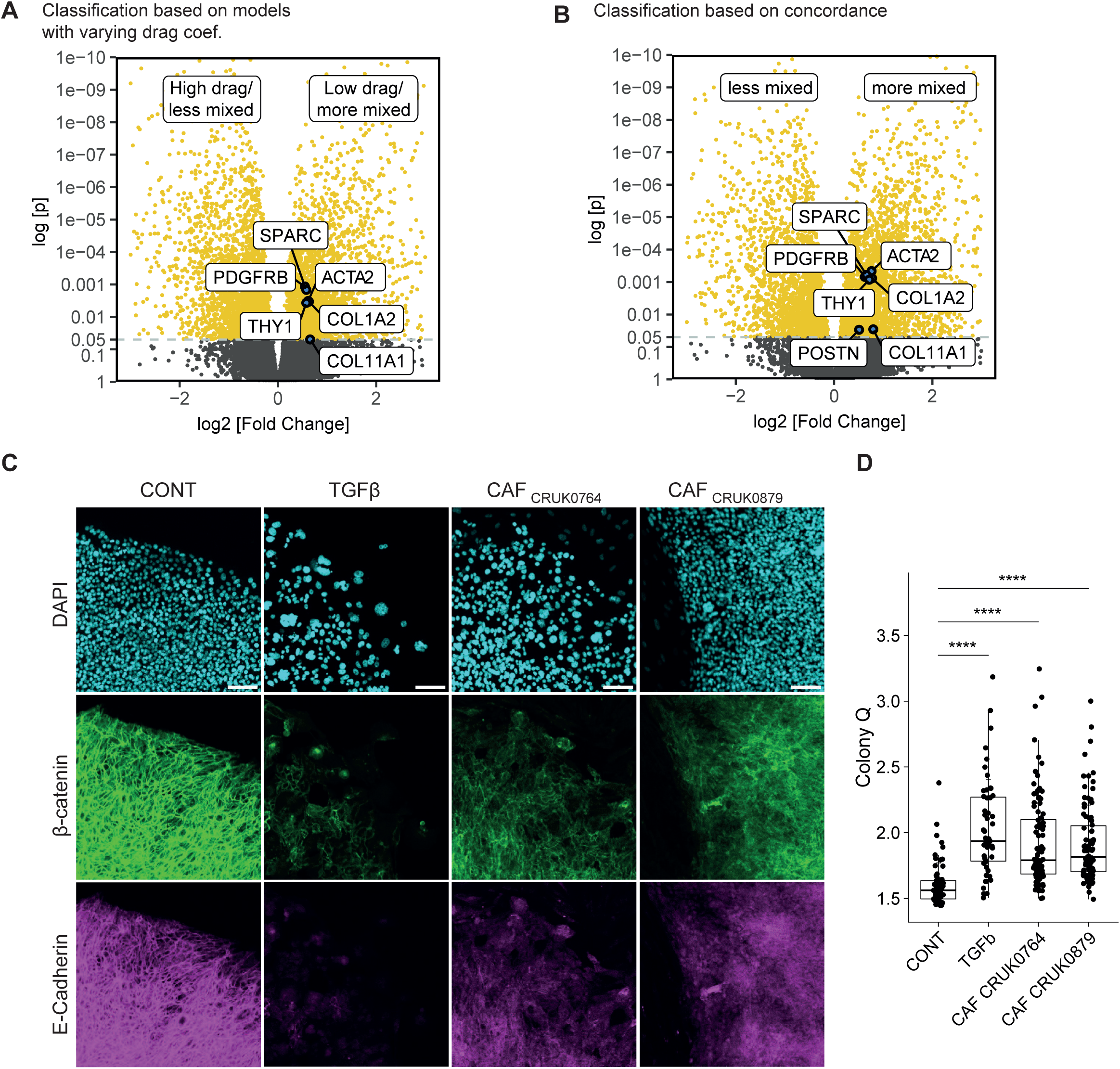
Cancer associated fibroblasts modulate subclonal mixing. A. Differential expression of genes between subsets of tumours inferred to be “more mixed” (N = 61 regions from 20 tumours) and “less mixed” (N = 60 regions from 20 tumours). Select significantly differentially expressed CAF and ECM marker genes are highlighted in the plot. Inference of mixedness was based on a SVM classifier trained using simulations with low vs high levels of drag coefficient. B. Differential expression of genes between subsets of tumours inferred to be “more mixed” (N = 61 regions from 20 tumours) and “less mixed” (N = 47 regions from 15 tumours). Select significantly differentially expressed CAF and ECM marker genes are highlighted in the plot. Inference of mixedness was based on concordant classifications between motility- and drag-based SVM classifiers. C. Representative immunofluorescence staining of DAPI, β-catenin and E-Cadherin and in KP control, TGFβ treated and KP-CRUK0764 and KP-CRUK0879 co-culture single cell colonies at the boundary. Scale bar = 100 μm. D. Quantification of colony Q = (perimeter / √ area) / π for Control n = 65, TGFβ n = 51, KP-CRUK0764 n = 88, KP-CRUK0879 n = 90 colonies per condition, N = 4 independent experiments. Error bars are mean ± sd, pairwise comparisons by Dunn’s test with Benjamin-Hochberg p-value correction, **** p < 0.0001. In (D), two ends of the box reflect the lower and upper quartiles, respectively, and the horizontal line dividing the box reflects the median. The red dot and error bar indicate the mean and the standard deviation, respectively.

**Supp. Figure 8:**
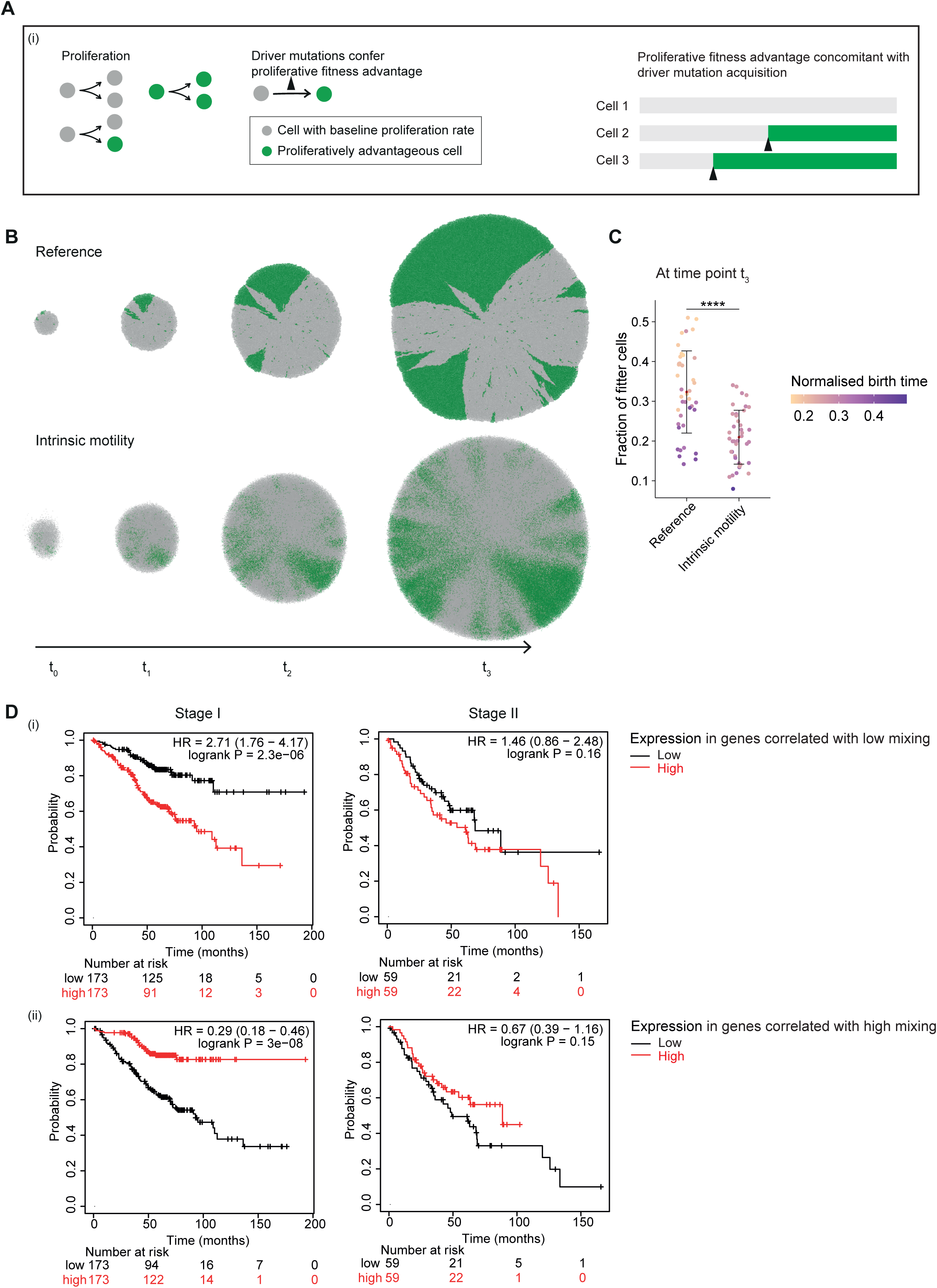
Computational modelling links subclonal mixing with differential outgrowth rates of fitter subclones. A. Schematics illustrating model implementation of proliferative fitness advantage. Proliferatively advantageous cells (green) emerge from acquiring a driver mutation upon proliferation. 5 out of 10000 alleles, when mutated, confer a proliferative advantage. Mutations of these alleles are referred to as driver mutations. B. Spatial patterns of proliferatively advantageous cells (green) over time in representative simulated tumours with low and high random cell motility, respectively. C. Fraction of fitter cells in proliferation at a late stage of simulated tumours with low and high random cell motility, respectively. N = 40 simulations for each condition. Data points are colour-coded according to the normalised birthtime of the largest subclone in the respective simulation. Welch two-sample t-test is performed. “****” reflects p <0.0001. The t-value and degree of freedom, (t, df), is: (5.8188, 67.346). D. Kaplan Meier plots shown the relationship between gene expression linked to low or high levels of inferred subclone mixing and patient outcomes. Upper panels show the association between genes correlated with low levels of mixing and overall survival in stage I and stage II lung adenocarcinoma. Lower panels show the association between genes correlated with high levels of mixing and overall survival in stage I and stage II lung adenocarcinoma. In (C), two ends of the box reflect the lower and upper quartiles, respectively, and the horizontal line dividing the box reflects the median. The red dot and error bar indicate the mean and the standard deviation, respectively.

